# The anti-neural role of BMP signaling is a side effect of its global function in dorsoventral patterning

**DOI:** 10.1101/2025.06.08.658475

**Authors:** Paul Knabl, June F. Ordoñez, Juan Daniel Montenegro Cabrera, Tim Wollesen, Grigory Genikhovich

**Affiliations:** Department of Neurosciences and Developmental Biology, University of Vienna, Austria; Vienna Doctoral School of Ecology and Evolution (VDSEE), University of Vienna, Austria; Konrad Lorenz Institute for Evolution and Cognition Research, Klosterneuburg, Austria; Department of Evolutionary Biology, University of Vienna, Austria

## Abstract

In Bilateria with centralized nervous systems (e.g. in vertebrates or arthropods), the minimum of the BMP signaling activity gradient defines the position of the central nervous system. BMP-dependent patterning of the secondary body axis is ancestral for Bilateria and possibly also for the bilaterian sister clade Cnidaria. However, the variety of levels of centralization of the nervous systems in Bilateria – from diffuse to fully centralized – as well as the lack of centralization of the nervous system in Cnidaria, suggest that BMP signaling cannot be perceived as a universally “anti-neural” signal. Here we use transgenic reporter lines in the anthozoan cnidarian *Nematostella* to show that BMP signaling is active in distinct neuronal populations. Moreover, attenuation of BMP signaling followed by RNA-Seq shows that BMP signaling is a positive regulator of many neuronal genes, including the top-tier neural progenitor marker *soxB(2)*. Further, we analyze BMP signaling activity in the true jellyfish *Aurelia* and box jellyfish *Tripedalia* proving that BMP signaling in the diffuse cnidarian nervous system is not an anthozoan but an ancestral cnidarian feature, shared by anthozoans and medusozoans. Finally, we show that the highly centralized ventral nervous system of the non-model spiralian, the chaetognath *Spadella*, forms out of paired BMP signaling-positive domains on the lateral sides of the embryo. Together, our data suggest that one of the ancestral roles of BMP signaling was in promoting neurogenesis. We propose that the “anti-neural” function of BMP signaling in vertebrates and arthropods is a side effect of its global role in the dorsoventral patterning of the ectoderm.

## Introduction

The evolutionary origin of BMP signaling-dependent regulation of animal nervous systems is uncertain. Both in vertebrates and arthropods, the establishment of a BMP signaling gradient is required for patterning the dorsoventral (DV) body axis and the formation of the centralized nervous system (CNS). In these models, the position of the neuroectoderm is defined by the inhibition of BMP signaling on one side of the DV axis – dorsally in vertebrates and ventrally in arthropods (Mizutani & Bier, 2008). Later during development, similar genetic programs regulate the medio-lateral patterning of the vertebrate neural plate and the nerve cords of insects and annelid worms (Denes et al., 2007; Sprecher & Reichert, 2003). These observations made in the mouse, frog, zebrafish, fly, and ragworm have contributed to the idea that the CNS emerged once, early during bilaterian evolution on the side of the body, where BMP signaling was low (Arendt et al., 2008; Arendt & Nubler-Jung, 1996; Arendt et al., 2016; De Robertis, 2008; Lichtneckert & Reichert, 2005). DV axis inversion at the base of chordates made this ancestrally ventral BMP minimum dorsal in chordates (Su et al., 2019).

While the role of BMP signaling in regulating the DV patterning is clearly ancestral and, with few exceptions, present across all Bilateria (Mörsdorf et al., 2024), its strictly “anti-neural” function is contradicted by the variety of the degrees of centralization of the bilaterian nervous systems – from diffuse to being arranged into a varying number of nerve cords to fully centralized (Martin-Duran & Hejnol, 2021; Moroz, 2009). Similarly variable are the effects of BMP signaling perturbations on the nervous systems of different groups, from promoting to suppressing neurogenesis to having no effect at all (Lowe et al., 2006; Lu et al., 2012; Lyons et al., 2020; Martin-Duran et al., 2018; McClay et al., 2018; Webster et al., 2021). Clearly, in order to understand the ancestral role of BMP signaling in neuron formation we need a wider phylogenetic sampling within and outside Bilateria, especially in animals with diffuse nervous systems and with centralized nervous systems forming in “non-standard” locations.

The evolutionary sister clade to Bilateria is the phylum Cnidaria, encompassing Anthozoa (sea anemones and corals) and four classes of Medusozoa: Hydrozoa (hydroids), Staurozoa (stalked jellyfish), Scyphozoa (true jellyfish) and Cubozoa (box jellyfish) (**Fig 1A**). Cnidaria possess a full repertoire of BMP pathway genes, including BMP ligands, BMP antagonists, BMP receptors and BMP effectors (Genikhovich & Technau, 2017; Gold et al., 2019; Kayal et al., 2018; Khalturin et al., 2019; Leclère et al., 2019; Lewis Ames et al., 2016; Ohdera et al., 2019). Cnidarian nervous systems are organized as diffuse nerve nets with local condensations, forming nerve rings and neurite tracts, but lacking any brain-like centralization (Havrilak et al., 2017; Marlow et al., 2009; Nakanishi et al., 2012). Neurons are found both in the epidermis and in the gastrodermis of cnidarians, and, strikingly, work on the anthozoan sea anemone *Nematostella vectensis* showed that gastrodermal neurons are “born” in the gastrodermis rather the migrate there, as is the case in Bilateria (Nakanishi et al., 2012). Neuronal gene expression in *Nematostella* commences already in the blastula, which is before pSMAD1/5 becomes first detectable (Knabl et al., 2024; Rentzsch et al., 2017; Watanabe et al., 2014). Neuronal markers were not affected by the disruption of BMP signaling during early phases of neurogenesis (Watanabe et al., 2014). Only at planula larva stage, the upregulation and, surprisingly, also downregulation of BMP signaling both reduced neuronal markers and the number of RFamide- and GLWamide-positive neurons (Watanabe et al., 2014) suggesting that neural “induction” is independent of BMP signals, while later neurogenesis may involve a combination of BMP activation and repression. Taken together, it is largely unclear, if and to which extend BMP signals are required during cnidarian neurogenesis.

**Fig 1.**
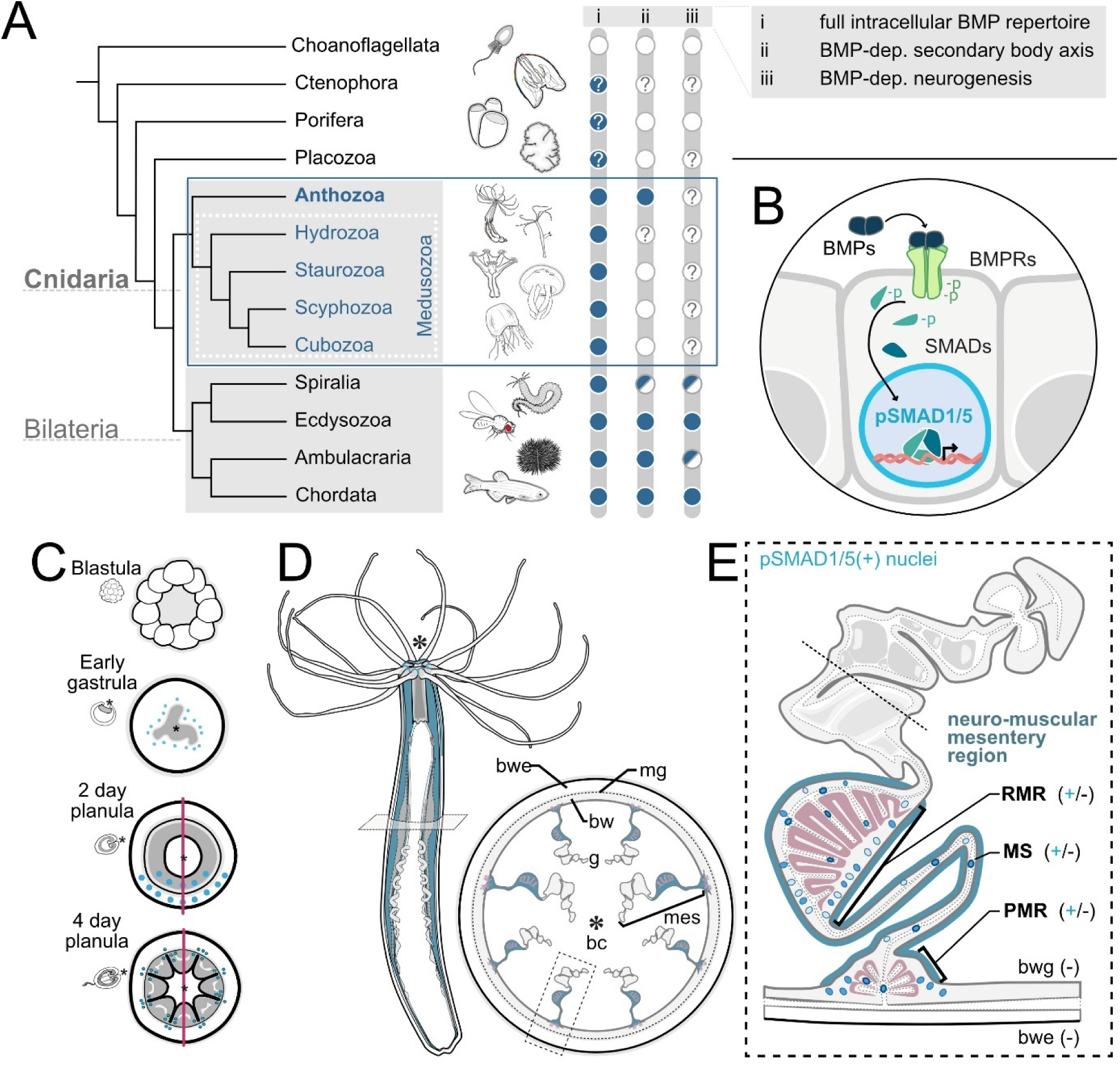
BMP signaling in *Nematostella vectensis*. (A) Simplified phylogenetic tree of early branching metazoans indicating the presence or absence of (i) BMP pathway genes, (ii) the presence of a BMP-dependent secondary body axis, (iii) Involvement of BMP signaling in neurogenesis. (B) Schematic of the BMP signaling pathway, BMP signaling is activated by the binding of BMP dimers to the BMP receptor complex, resulting in the nuclear translocation of phosphorylated SMAD1/5 (pSMAD1/5), acting as BMP effector and regulating BMP-responsive gene expression. (C) BMP signaling dynamics during early development of *Nematostella*, no activity in the blastula, pSMAD1/5 activity around the blastopore in the early gastrula, pSMAD1/5 gradient at 2-day planula stage and dispersed pSMAD1/5 activity in the 4-day planula. (D) Overview of *Nematostella* polyp anatomy in lateral view and as a cross-section of the body column, the neuromuscular domain of the mesentery gastrodermis is highlighted in blue. (E) Detailed schematic of an individual mesentery as a cross-section, highlighting the localization of pSMAD1/5-positive nuclei within the neuro-muscular domain.

Most of our knowledge about BMP signaling in Cnidaria comes from an anthozoan model – the sea anemone *Nematostella vectensis*. Unlike radially symmetric Medusozoa, *Nematostella* and other Anthozoa, display a bilaterally symmetric body plan with a secondary, “directive” body axis (**Fig 1A**). Similar to the bilaterian DV axis, the directive axis requires the formation of a BMP signaling gradient that is regulated by the interplay of multiple BMP pathway components, including *bmp2/4, bmp5-8, gdf5-like* (=*gdf5-l*)*, chordin, gremlin*, *rgm* and *zswim4-6* (Genikhovich et al., 2015; Knabl et al., 2024; Leclère & Rentzsch, 2014; Saina et al., 2009). Active BMP signaling can be visualized by the spatial distribution of its transcriptional effector, phosphorylated SMAD1/5 (pSMAD1/5). In *Nematostella*, at late gastrula stage, BMP signaling forms a pSMAD1/5 gradient along the directive axis. This gradient persists while the directive axis is being patterned, but eventually disappears by 4 day planula larva stage, at which point BMP signaling activity becomes local and especially prominent in the mesenteries – the gastrodermal septa of the bodywall (Genikhovich et al., 2015; Knabl et al., 2024; Leclère & Rentzsch, 2014). Our recent study revealed pronounced BMP signaling activity in the mesenteries of the adult *Nematostella* polyp (Knabl et al., 2025). The mesenteries harbor a diversity of cell populations including epithelial, digestive, reproductive, muscular and neuronal cells. The two latter are concentrated in the “neuro-muscular” mesentery region, where the longitudinal muscles (the retractor and parietal muscle), the longitudinal neurite tracts, and sensory and ganglion mesentery neurons are located. BMP signaling is most pronounced within the neuro-muscular domain, yet it was unclear if BMP signaling occurs in neuronal or muscular cell populations (**Fig 1**). Here, we analyze the potential involvement of BMP signaling in the diffuse nervous system of the anthozoan cnidarian *Nematostella*. Additionally, we use two other distantly related medusozoan cnidarians *Aurelia* and *Tripedalia* for comparison and show a pro-neural action of BMP signaling in these models. Finally, we also demonstrate distinct BMP activity in the central nervous system of the non-model spiralian *Spadella*. Together, our findings suggest that BMP signaling was ancestrally a pro-neural factor.

## Results

### BMP pathway genes are expressed across neuronal subtypes in the developmental single-cell atlas of *Nematostella*

To address the expression of BMP components in the neuroglandular and muscular repertoire of *Nematostella*, we utilized the updated developmental single-cell atlas of *Nematostella* (Cole et al., 2024). The analysis of the neuroglandular subset by Cole et al. presents 47 developmentally closely related but distinct cell states, including *insulinoma (insm)*-positive (N1) and insm-negative (N2) neurons, secretory cells (S) and digestive gland cells (GD). Among the N1 and N2 neurons, several gastrodermis-derived types have been identified so-far, including six N1 (N1.g1-6) and a single N2 type (N2.g). The analysis of the retractor muscle subset by Cole et al. revealed four distinct cell states, including the early retractor muscle state (RM.1), the late tentacle retractor muscle state (TR.2), the early tentacle retractor muscle state (TR.1), and the late retractor muscle state (MR.1). We generated expression plots of BMP pathway genes, including BMP ligands (*bmp2/4, bmp5-8, gdf5-l, admp*), BMP effectors (*smad1/5, smad4, smad4-like*), BMP receptors (*alk2, alk3/6, bmprII, actrII*) and intracellular, membrane-bound, and secreted BMP antagonists (*rgm, smad6, chordin, gremlinA, gremlinB, noggin1, noggin2, crosveinless2 and follistatin*) (Knabl et al., 2025). The expression of BMP components was detectable in N1 and N2 neuron types, while it was reduced in putative sensory neurons (N1S) and secretory cells (S) and absent in digestive gland (GD) types (**Fig 2A**). Notably, the expression of BMP components in N1 and N2 neurons is not limited to the gastrodermal populations, despite the fact that our previous analysis revealed that BMP signaling activity is largely limited to the mesentery gastrodermis (Knabl et al., 2025). The expression of extracellular molecules such as BMP ligands and antagonists appeared to be sparse with only small fractions of cells (<10%) in each subpopulation expressing the transcripts (**Fig 2A**). Moreover, single-cell analysis suggested that neither BMP pathway genes nor BMP receptor type expression is specific to discrete neuronal subpopulations. While our expression analysis of the developmental single-cell data pointed to the expression of the BMP module in neuronal populations, it was not sufficient to identify specific candidates among the neuronal subpopulations, which might display pSMAD1/5 activity. The expression of BMP pathway genes in the retractor muscle cell states was low or absent, except the upregulated expression of BMP ligands in the early tentacle retractor muscle TR.1 and of the BMP antagonist molecule RGM in the late tentacle retractor muscle state TR.2 (**Fig 2B**).

**Fig 2.**
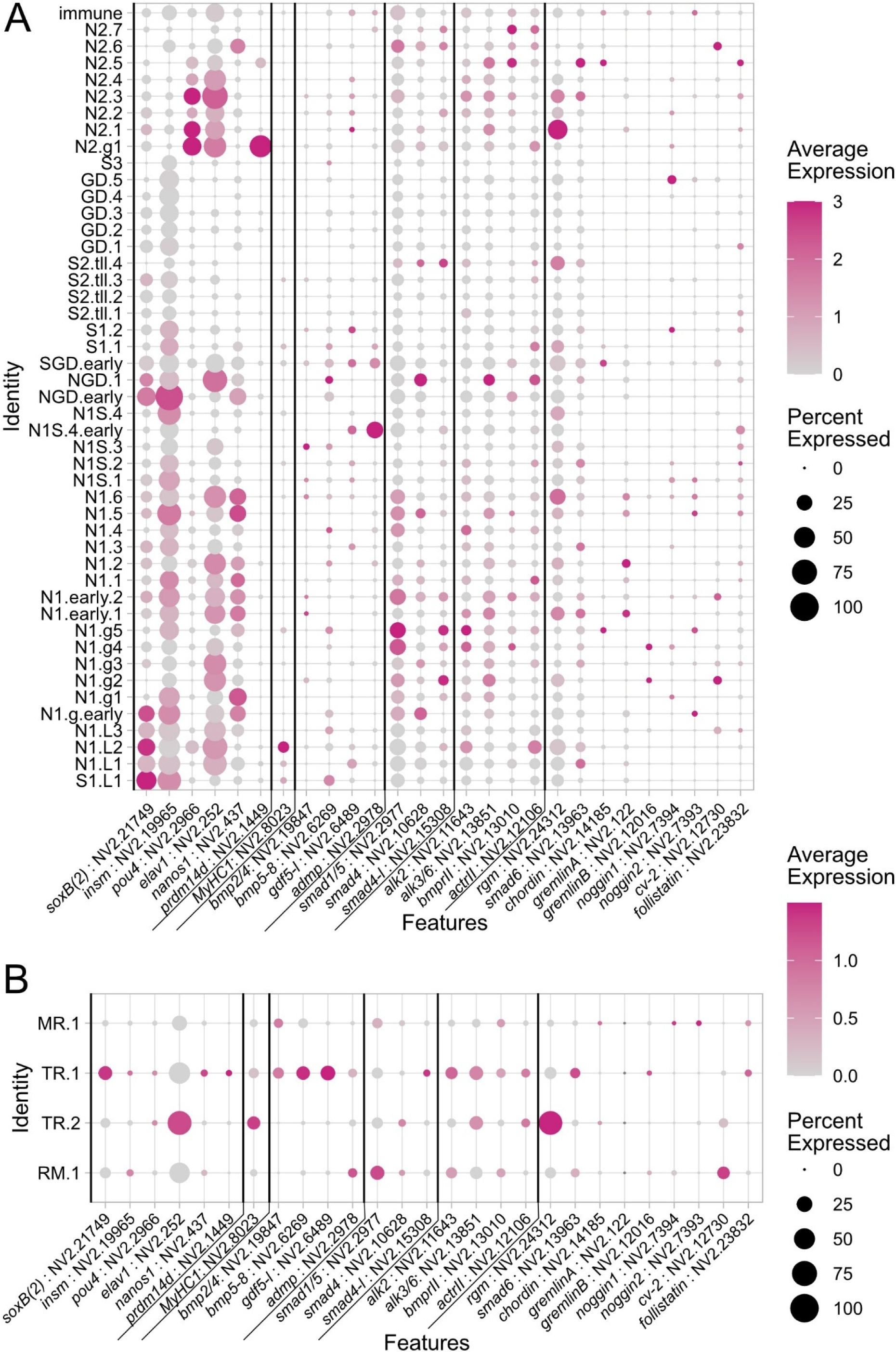
Expression of BMP pathway genes in the neuronal and retractor muscle subset of the developmental single-cell atlas of *Nematostella*. Dot plots showing the expression of neuronal marker genes, retractor muscle marker *myosin heavy chain1* (*myhc1*) and BMP signaling components (A) in the neuroglandular subset and the (B) retractor muscle subset of the *Nematostella* developmental single cell by Cole et al. 2024. Average scaled expression of 0 or below is indicated in grey.

### pSMAD1/5 is active in the neurons but not in the muscle cells of the neuro-muscular region of the mesentery

Given the upregulated expression of BMP pathway genes in neuron types according to our single-cell analysis, we aimed to determine if BMP signaling is indeed active in neural cell populations using different transgenic reporter lines. Earlier, we showed that BMP signaling activity in the adult polyp is most pronounced in the neuro-muscular region of the mesentery gastrodermis (Knabl et al., 2025), therefore, we focused our analysis on reporter lines with expression in gastrodermal neurons and muscles. In the neuro-muscular region, accumulations of different neurons are found along the retractor and the parietal muscles as well as in the mesentery stalk (**Fig 1E**, **Fig 3**). Firstly, we examined the retractor muscle region, containing the retractor muscle and multiple ganglionic and sensory neuron populations. The expression plot of the single-cell retractor muscle subset suggests low expression levels of BMP components in the retractor muscle (**Fig 2B**). To determine if BMP signaling is active in the retractor muscle cells, we performed anti-pSMAD1/5 antibody staining in the *myosin heavy chain1::mCherry* (*myhc1::mCh*) transgenic reporter line, expressing mCherry under the *myhc1* promotor in the retractor muscles in the mesenteries as well as in the epidermal muscles of the tentacles (Renfer et al., 2010). In *myhc1::mCh* transgenic animals, we observed pSMAD1/5-positive cells occurring within the gastrodermal epithelium or within the mesoglea as basoepithelial cells (**Fig 3B-C**). These pSMAD1/5-positive cells, however, displayed no overlap with mCherry-positive retractor muscle cells (**Fig 3C-F’**), suggesting BMP signaling is not active in the retractor muscle.

**Fig 3.**
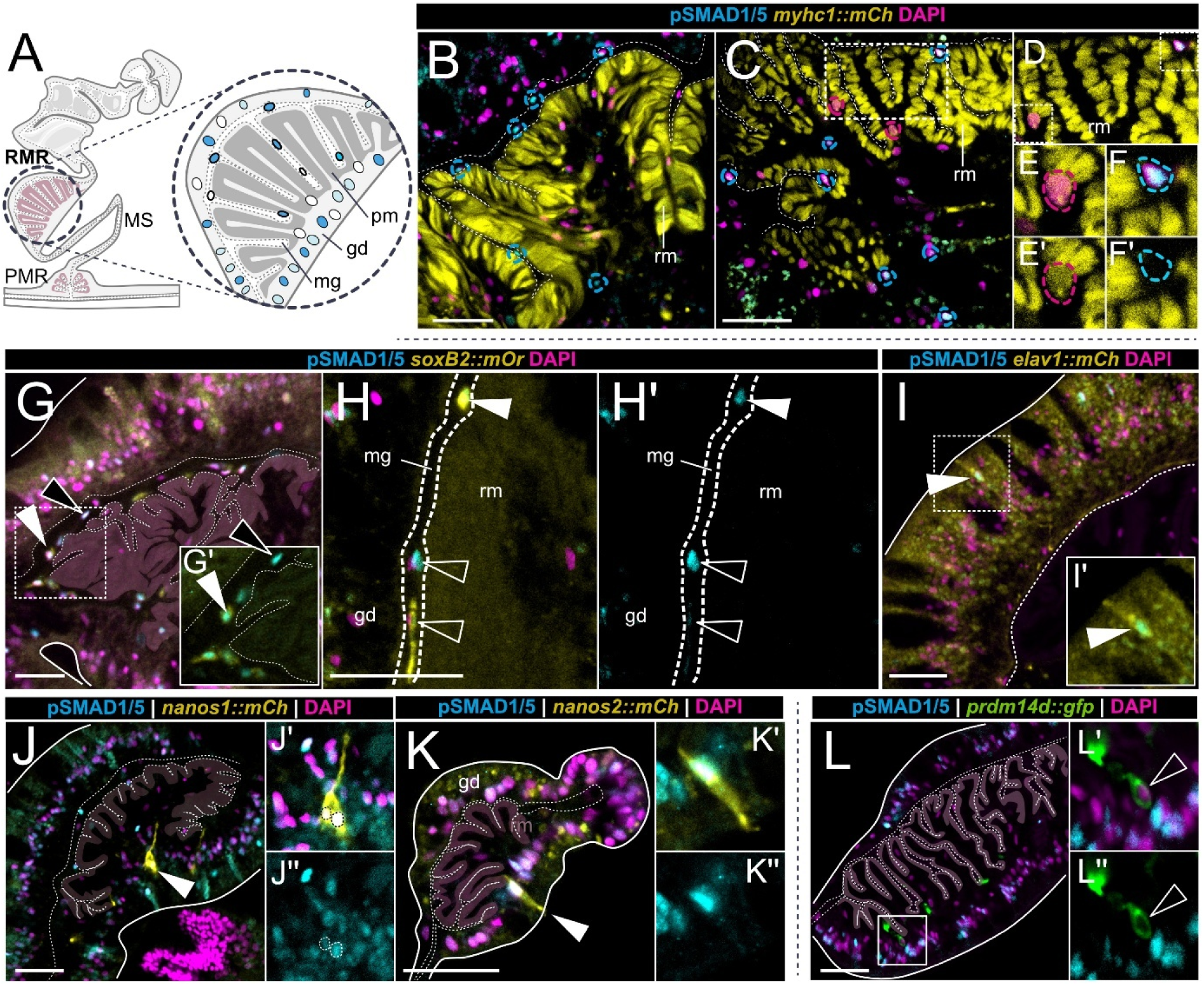
BMP signaling is active in specific neuron types located in the retractor muscle region of the mesentery. (A) Schematic overview of a mesentery cross-section, highlighting the retractor muscle region (RMR) within the neuromuscular domain. (B-K) Immunostaining on cross-sections of the retractor muscle region for pSMAD1/5 (cyan) and nuclei (magenta) in transgenic reporter lines. (B-F) *myosin heavychain 1* reporter expressing mCherry (yellow) (*myhc1::mCh*) showing pSMAD1/5-positive nuclei located in the mesoglea and gastrodermis, (C-F) detail, showing no overlap between pSMAD1/5-positive nuclei (cyan dashed outlines) and mCherry-positive nuclei of retractor muscle cells (magenta dashed outlines). (G-H) pSMAD1/5-positive nuclei overlap (white arrowhead) with mOrange in the *soxB(2)* reporter line (*soxB(2)::mOr*), (I) pSMAD1/5 activity overlaps with mCherry-positive cells in the *elav1* reporter line (white arrowhead). (J) *nanos1* reporter expressing mCherry (*nanos1:mCh*) shows pSMAD1/5 staining in *nanos1*-positive putative ganglionic cells (white arrowhead), (K) *nanos2* reporter expressing mCherry (*nanos2::mCh*) shows pSMAD1/5 staining in the *nanos2*-positive putative sensory cells (white arrowhead), (L) *prdm14d* reporter expressing GFP (*prdm14d::gfp*) does not show pSMAD1/5 staining in the *prdm14d*-positive cells (white outlined arrowhead). immunostaining for pSMAD1/5 (cyan), DAPI (magenta), reporter protein (yellow or green). Scale bars 25µm. White arrow heads indicate overlap, white outlined arrowheads indicate no overlap. gd–gastrodermis, pm–parietal muscle, mg–mesoglea, ep–epidermis.

Next, we tested if BMP signaling is active in neuronal subpopulations with known localization in the neuro-muscular mesentery, including *soxB(2)*, *elav1*, *nanos1* and *prdm14d* expressing cells (Denner et al., 2024; Lemaitre et al., 2023; Miramon-Puertolas et al., 2024; Nakanishi et al., 2012; Richards & Rentzsch, 2014; Steger et al., 2022). The *soxB(2)::mOr* reporter line was previously shown to mark neuronal progenitors and differentiated neurons in the regions of the retractor muscle, parietal muscle and body wall gastrodermis (Miramon-Puertolas et al., 2024). We observed a partial overlap of *soxB(2)* expression and pSMAD1/5 activity in cells of the retractor muscle region, primarily located within the mesoglea (**Fig 3G**, white arrowheads). These basoepithelial cells were mostly pSMAD1/5- and *soxB(2)*-double-positive, while we rarely observed only pSMAD1/5-positive or only *soxB2*-positive cells, suggesting that a large portion of neural progenitor cells of the mesentery receives BMP signals (**Fig 3G-H’**). We then examined transgenic reporter lines for *elav1* and *nanos1*, marking sensory and ganglionic neurons in the gastrodermis and epidermis (Nakanishi et al., 2012; Steger et al., 2022) and another *nanos* paralog, *nanos2*, which labels a broad range of cell types in both body layers, including neuroglandular populations in the mesentery (Denner et al., 2024). In *elav1::mCherry*, *nanos1::mCherry* and *nanos2::mCherry,* we discerned an overlap of pSMAD1/5 with neuronal populations (**Fig 3I-K’’**, white arrowheads). For *nanos1,* double-positive mCherry and pSMAD1/5 cells had a ganglionic neuron morphology (**Fig 3J-J’’**), while for *elav1* and *nanos2*, we found double-positive cells, which looked like sensory neurons (**Fig 3I-I’,K-K’’**). The situation was different when we analyzed the *prdm14d::gfp* reporter line, in which GFP marks ganglionic neurons in the mesentery retractor and parietal muscle region (Lemaitre et al., 2023). In the *prdm14d::gfp* line, GFP-positive cells never displayed BMP signaling activity (**Fig 3L-L’’**), suggesting BMP signaling in the mesentery retractor region is activated only in distinct neuronal subtypes.

Next, we investigated the basal-most domain of the neuromuscular mesentery, where the parietal muscle region is located. Here, pSMAD1/5-positive cells form two gastrodermal clusters that are ambilateral to the parietal muscles and follow the entire primary body axis length (**Fig 4**). The same area is known to contain the longitudinal neural tracts, two parallel bundles of concentrated neurites on each side of the parietal muscle. To check if BMP signaling coincides with neurons in the parietal muscle region, we analyzed pSMAD1/5 immunoreactivity in the reporter lines for *soxB(2)*, *elav1, nanos1* and *prdm14d,* which have been demonstrated to label the neural tracts (Havrilak et al., 2017; Layden et al., 2016; Marlow et al., 2009; Nakanishi et al., 2012; Rentzsch et al., 2017; Steger et al., 2022). First, we tested if BMP signaling is active in *soxB2::mOr* cells and detected pSMAD1/5 in mOrange-positive basiepithelial cells in the mesoglea of the mesentery stalk and parietal muscle region (**Fig 4B-C’**, white arrowheads). In addition, BMP signaling was active in *soxB2::mOr*-positive cells with differentiated sensory neuron or ganglionic neuron morphology, which were embedded in the epithelium of the parietal muscle region (**Fig 4D-D’**, white arrowheads). To further examine differentiated neuronal types, we analyzed the parietal muscle region in the reporter lines *elav1::mCh* and *nanos1::mCh* (Nakanishi et al., 2012; Steger et al., 2022). Both reporter lines label sensory and ganglionic neurons in both body layers, with *elav1::mCh* expression being especially pronounced in the neurite tracts (Nakanishi et al., 2012; Steger et al., 2022). In the *elav1::mCh* and *nanos1::mCh* reporter line, we discerned cells with sensory neuron-like appearance that were pSMAD1/5-positive (**Fig 4E-F’’**, white arrowheads). Notably, pSMAD1/5 was again present in cells not expressing fluorescent reporter proteins, suggesting the presence of other cell populations with active BMP signaling along the parietal muscle region (**Fig 4E-F)**. Similar to the situation in the retractor muscle region, the *prdm14d::gfp* reporter line, which labels ganglionic neurons along the neurite tracts (Lemaitre et al., 2023), showed no pSMAD1/5 in the *prdm14d::gfp-*positive cells in the parietal muscle region (**Fig 4G-I’**, white outlined arrowhead), similar to the situation in the retractor muscle region (**Fig 3L**). These findings suggest that BMP signaling is active in specific parts of the gastrodermal nervous system, where it is restricted to neuronal progenitors and distinct ganglion and sensory neurons of the mesentery, while it is absent from distinct subpopulations of the neuronal lineage in the same regions.

**Fig 4.**
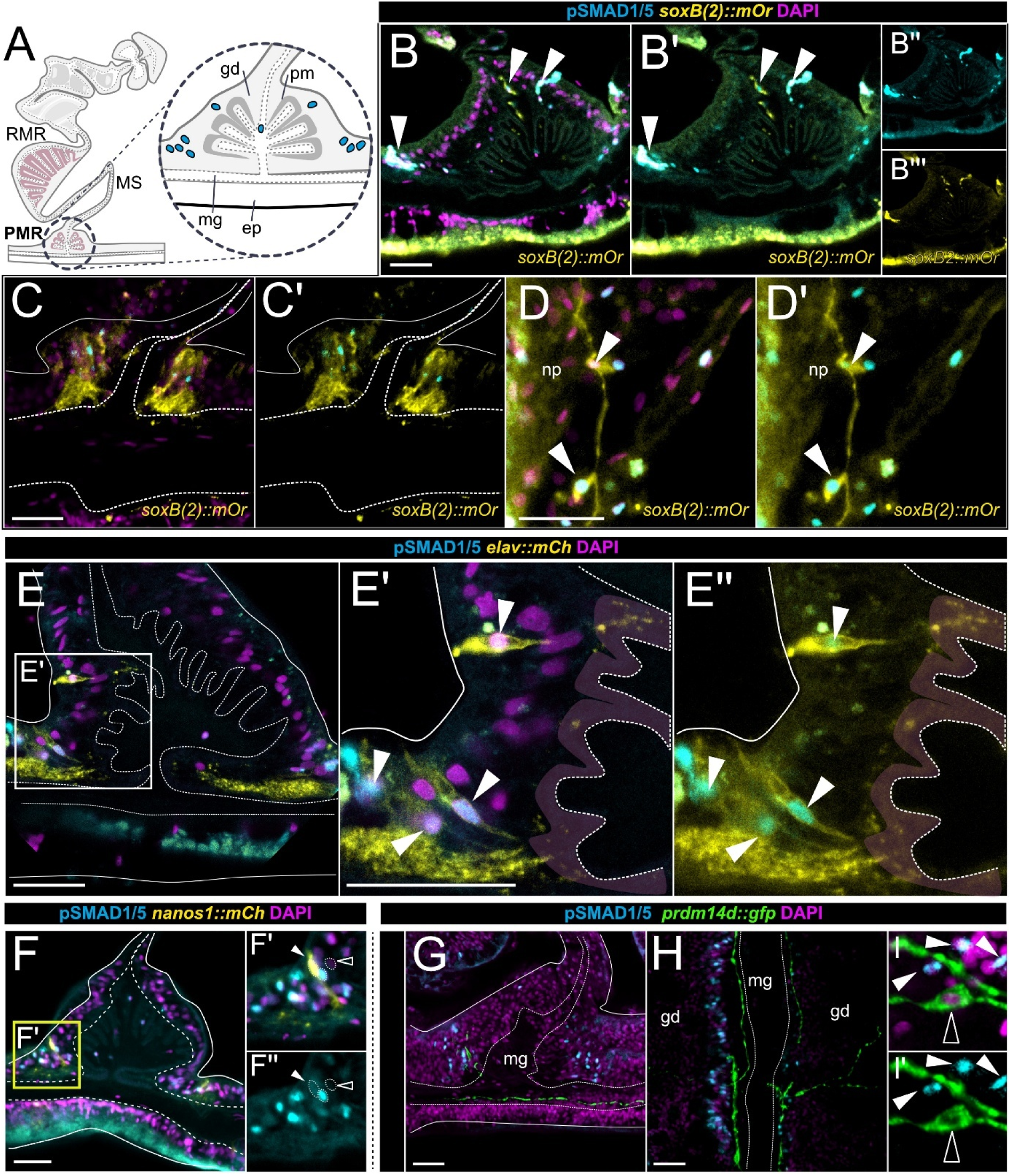
BMP signaling is active in specific neuron types of the parietal muscle region of the mesentery. (A) Schematic overview of a mesentery cross-section, highlighting the parietal muscle region (PMR) within the neuromuscular domain. (B-H) Immunostaining for pSMAD1/5 (cyan) and nuclei (magenta) in the parietal muscle region of transgenic reporter lines as cross-sections or as lateral views. (B-D) mCherry-positive cells (yellow) in the *soxB(2)* reporter line (*soxB(2)::mCh*), located in the mesoglea and gastrodermis, partially overlap with pSMAD1/5 activity (white arrowheads), (B-C) in cross-sections and (D) lateral view. White arrowheads in (E-F) indicate pSMAD1/5-positive nuclei (cyan) in the epithelium of the parietal muscle region overlap (E) with mCherry (yellow) in the *elav1* reporter line (*elav1::mCh*) and (F) with mCherry (yellow) in the *nanos1* reporter line(*nanos1::mCh*). (K) No overlap (white outlined arrowhead) of pSMAD1/5 (cyan) and GFP (green) detectable in the *prdm14d* reporter line (*prmd14d::gfp*). gd – gastrodermis, pm – parietal muscle, mg – mesoglea, ep – epidermis.

### BMP signaling attenuation in the adult *Nematostella* polyp results in the differential expression of neuronal regulators

To better understand the BMP signaling-dependent gene regulation in adult *Nematostella*, we suppressed BMP signaling in the mature polyp using the BMP receptor inhibitor K02288 (**Fig 5A**), dissected the polyps into head without tentacles (from female polyps), body wall with mesentery base (from female polyps), and isolated mesenteries (from female and male polyps separately), and performed bulk RNA-Seq in biological triplicates. (**Fig 5B**). To enrich for direct rather than indirect downstream target genes, we wanted to keep inhibitor treatment as short as possible. We determined that 5 hours of K02288 treatment was the minimal duration sufficient to reliably inhibit BMP signaling in the polyp compared to the DMSO control as manifested by the disappearance of the nuclear pSMAD1/5 (**Fig 5A**). DeSeq2 analysis revealed effects of BMP inhibition in all tissues, with the differential expression (padj. of ≤ 0.05) of 203 genes in the head (145 down, 58 up), 233 genes in the body wall (206 down, 27 up), 56 genes in the female mesentery (53 down, 3 up) and 170 genes in the male mesentery (146 down, 24 up), comprising 489 unique genes differentially expressed across all tissues (**Fig 5C**). Among the different tissues only 9 genes were differentially regulated in all tissue samples (**Fig 5C, S1 Table**). To check for shared BMP signaling target genes across developmental stages, we compared the K02288/DMSO treatment dataset of the adult polyp with two recently published datasets of BMP signaling targets during early development: the dataset of direct BMP signaling targets at Late gastrula and 4 day planula stage identified by pSMAD1/5 ChIP-Seq, as well as the bulk RNA-seq dataset of the 2d planula upon BMP2/4 morpholino knockdown (bmp2/4MO) (Knabl et al., 2024). Out of 210 pSMAD1/5 ChIP target genes at gastrula and 4d planula (see Materials and methods for details), 20 genes were also differentially expressed upon K02288 treatment. Seven of these 20 were transcription factors or co-factors (**Fig 5D, S2 Table**). 289 genes were differentially expressed upon both K02288 treatment and bmp2/4MO knockdown, and 12 genes were present in all three datasets (**Fig 5E, S2 Table**).

**Fig 5.**
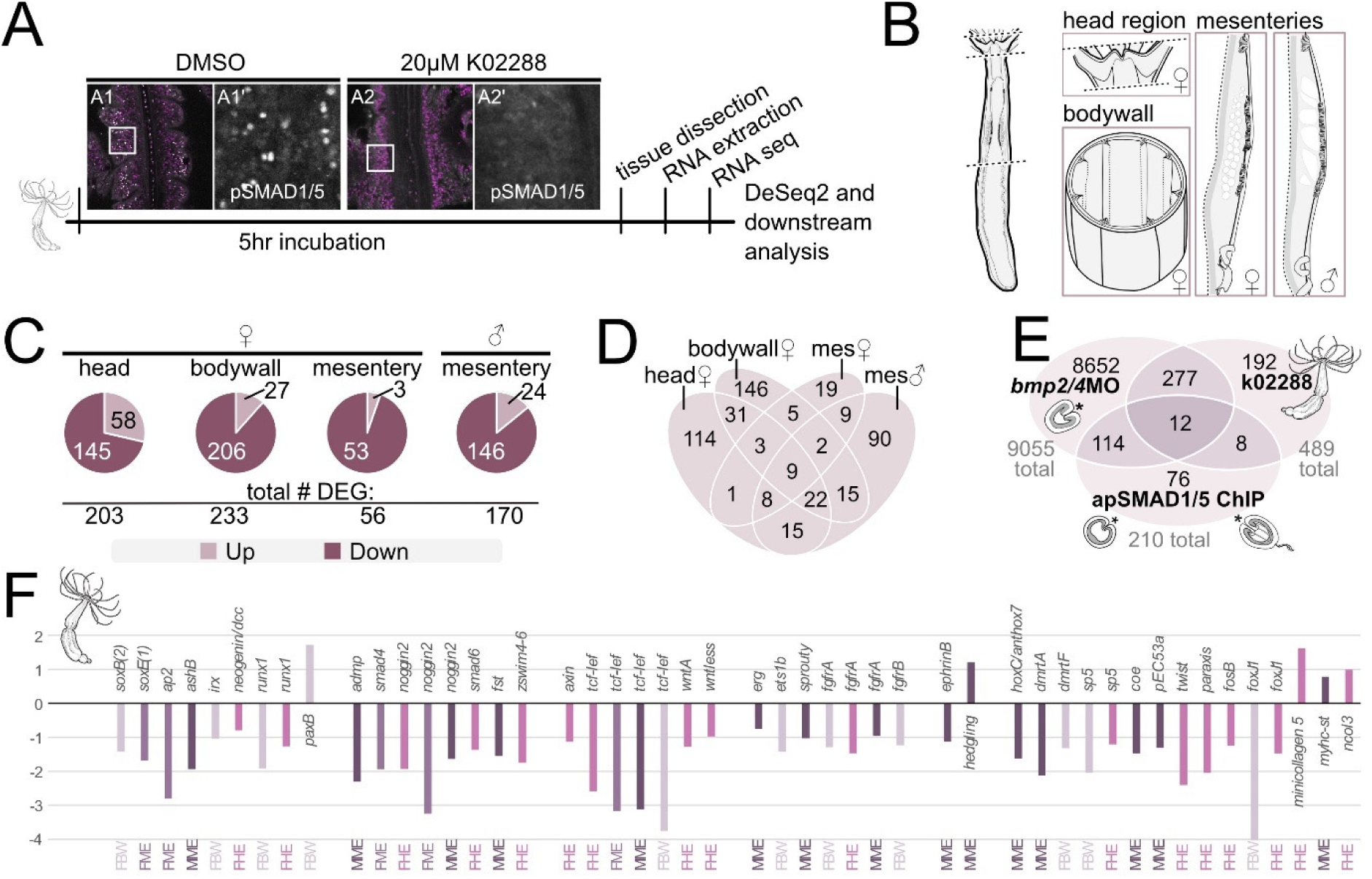
Pharmacological suppression of BMP signaling in the adult *Nematostella* polyp reduces the expression of developmental and neuronal regulators. (A) Schematic of BMP signaling inhibition of the adult polyp using K02288 treatment and downstream processing and analysis. (B) Schematic of tissue dissection for tissue-specific bulk RNA-seq (C) pie charts illustrating the number of down and upregulated genes upon K02288 treatment in individual tissues. (D) Venn diagram showing the relationships of (DE) differentially expressed genes between sets of tissues. (E) Venn diagram illustrating the relationships between the combined set of DE genes in K02288 treated adults and previously published data sets of DE genes in the 2-day planula upon bmp2/4 morpholino knockdown and direct pSMAD1/5 targets at Late gastrula and 4d planula (Knabl et al. 2024). (F) Bar plot showing log2 fold change (padj. < 0.05) of selected differentially expressed genes in adult polyp tissues (FBW-female body wall, FME-female mesentery, MME-male mesentery, FHE-female head region) upon K02288 treatment.

Both, in the RNA-seq datasets of BMP2/4 knockdowns and K02288 treatments, we identified various components of Wnt, FGF, TGFβ/BMP, Notch and MAPK signaling, of which some were also direct pSMAD1/5 target genes in *Nematostella* embryos (**S2 Table**). Previous works implicated multiple signaling pathways in nervous system development and maintenance in *Nematostella* (Rentzsch et al., 2017), however, we cannot treat differential expression of these “broad-spectrum” regulatory genes as evidence for or against the involvement of BMP signaling in neural development. In contrast, some of the differentially expressed genes were clearly neural factors, suggesting neuronal regulation by BMP signaling across developmental stages. Previous characterization of the direct pSMAD1/5 target genes in Late gastrula and 4-day planula stage and of the genes differentially expressed upon morpholino knockdown of BMP2/4 yielded multiple putative or verified neuronal genes (**S2 Table**) (Knabl et al., 2024). Similarly, the dataset of K02288 treatments of adult polyps shows an overrepresentation of the GO-terms for neurogenic processes among the differentially expressed genes: 332 of the 489 unique GO-terms are neurogenesis-related **(S1 Table, S1 File – Fig. 1).** The K02288 dataset of differentially expressed genes contains various putative or verified neuronal markers (*soxB(2), soxE1, ashB, paxB, ap2, runx, irx, dmbx, foxJ1, zswim4-6, ephrinB, c-ski,* etc.; **S1 Table**, **Fig 5F**), the majority of which are downregulated upon attenuation of the BMP signaling.

### Neural lineage markers are downregulated by BMP signaling attenuation in the *Nematostella* larvae

Previous analyses suggested that BMP signaling has both positive and negative effects on neurogenesis in the planula larvae (Watanabe et al., 2014). We searched for differentially expressed neural regulators in the bmp2/4MO RNA-seq data set, revealing multiple neuronal genes, most of which were significantly downregulated (**S2 Table**). We analyzed the expression patterns of selected neuronal genes upon the suppression of BMP signaling by BMP2/4 morpholino or K02288 treatment using in situ hybridization. First, we analyzed the expression of *soxC* and *soxB(2),* marking neuronal progenitors (Richards & Rentzsch, 2015; Steger et al., 2022), using bmp2/4MO knockdown, resulting in the complete loss of the pSMAD1/5 gradient, and gdf5-lMO knockdown, reducing graded pSMAD1/5 activity (Genikhovich et al., 2015). The expression of both *soxC* and *soxB(2)* was strongly reduced in bmp2/4MO (**Fig 6CD**). In contrast, *soxC* and *soxB(2)* were not affected by gdf5-lMO (**Fig 6D**), suggesting that either their expression requires input specifically from BMP2/4-mediated and not GDF5-l-mediated signaling or that reduced BMP signaling is sufficient to maintain their normal expression. Then, we analyzed the effect of the chemical inhibition of BMP signaling using the K02288 inhibitor, spanning different time windows: i) from Blastula (16-18 hpf) to 2d planula stage (**B->2dP**), overlapping with the time of the pSMAD1/5 gradient formation, ii) from 2d to 4d planula stage (**2dP->4dP**), when the pSMAD1/5 activity gradually switches from graded to dispersed and becomes located in the mesenteries, corresponding to the time span of gastrodermal patterning and mesentery formation, iii) from 4d to 5d (**4dP->5dP**), after the directive axis has been fully patterned and pSMAD1/5 is active in the mesenteries. For the different time windows of treatment, we again examined the expression of differentially expressed neural markers from the bmp2/4MO dataset, including *soxC* and *soxB(2),* as well as *insulinoma (insm),* which is expressed in neurosecretory and neuroglandular populations (Tourniere et al., 2022), *prdm14d* as marker for neural progenitors and ganglion neurons in the gastrodermis (Lemaitre et al., 2023) and *pou4* and *elav1* as neuronal differentiation markers of gastrodermal and epidermal subpopulations (Nakanishi et al., 2012; Tourniere et al., 2020). K02288 treatments from blastula to 2d planula resulted in strong downregulation of all analyzed marker genes (**Fig 6E**), concurring with the results of the bmp2/4MO in situ and RNA-seq data. Similarly, BMP inhibition starting at 2d planula until 4d planula stage, strongly reduced neuronal marker genes (**Fig 6E**). In comparison, K02288 treatments from the 4d to 5d planula, which is after the directive axis has been patterned, and mesenteries have been formed, displayed only a mild reduction of neural marker expression (**Fig 6E**), suggesting that the neuronal regulation becomes less sensitive to BMP signaling input in the fully patterned late planula. While early studies indicated a combination of positive and negative effects of BMP signaling on neural patterning in the planula, our data support a predominantly pro-neural influence of BMP signaling in the nervous system of the *Nematostella* larva.

**Fig 6.**
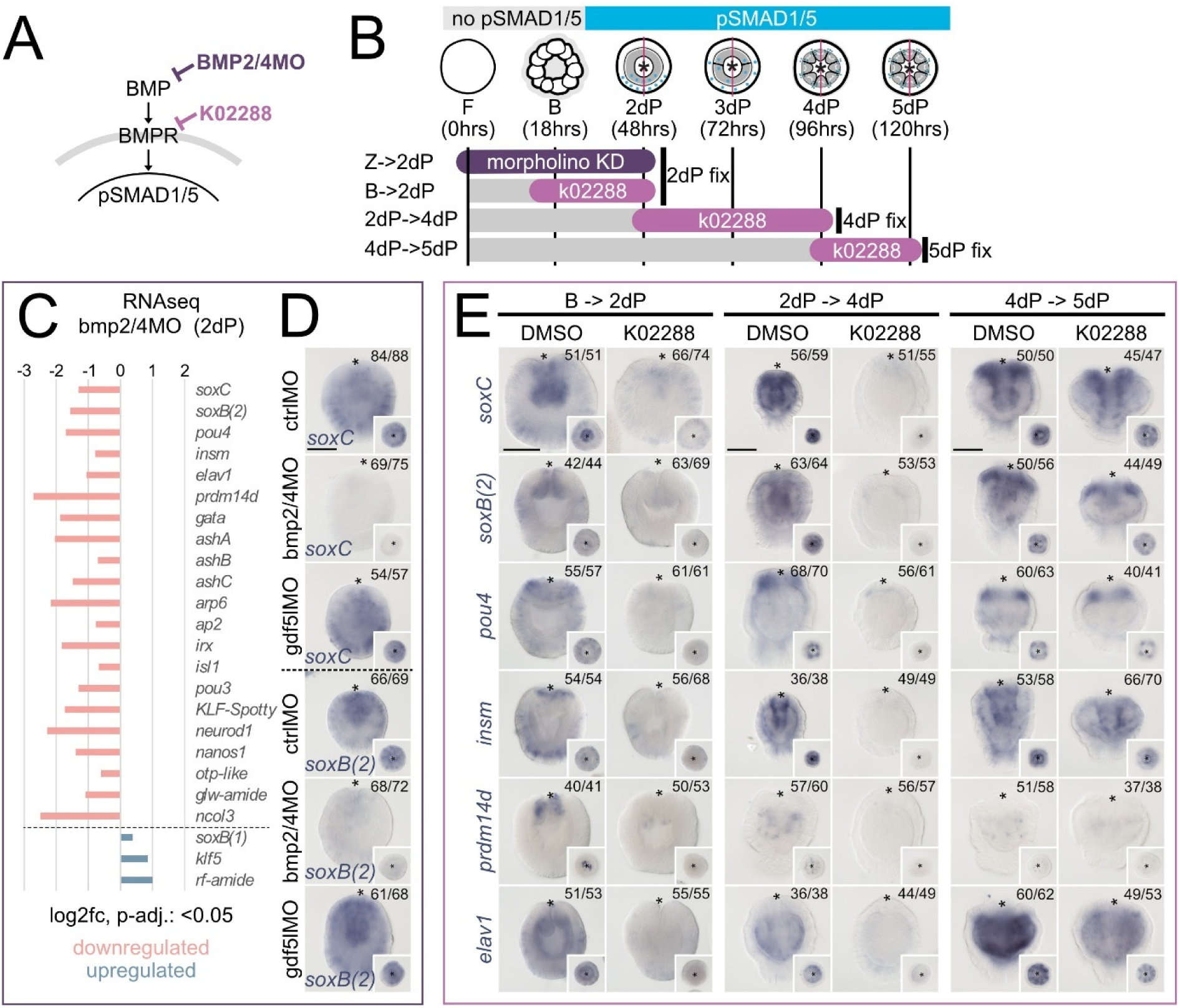
Neuronal markers are downregulated upon BMP signaling inhibition in *Nematostella* planula. (A) Schematic of the BMP signaling pathway and how it is affected by BMP2/4 morpholino oligo and pharmacological inhibition of the BMP receptor with K02288. (B) Timeline of *Nematostella* early development and BMP signaling dynamics from fertilization (F), Blastula (B) and 2-5 day Planula larvae. The grey/cyan-colored bars indicate the absence/presence of pSMAD1/5 activity. Duration and fixation time points of morpholino knockdown experiments (purple) and K02288 treatments (lilac). (C) Bar plot of expression changes (log2 fold, padj. <0.05;) of selected neuronal marker genes in 2 day planula upon bmp2/4MO knockdown by RNA-seq analysis (Knabl et al. 2024, see Methods). (D) Expression analysis of *soxC* and *soxB(2)* expression upon ctrlMO, bmp2/4MO, and gdf5-lMO knockdown at 2 day planula stage by in situ hybridization. (E) Expression analysis of selected neuronal marker expression upon K02288 treatments at 2 day planula, 4 day planula and 5 day planula stage by in situ hybridization. Asterisks indicate blastopore, scale bars 100µm.

### BMP signaling is active in the medusozoan nervous system

Life cycles of medusozoan cnidarians contain a pelagic medusa stage, whose behavior is much more complex than that of a sessile polyp, indicative of a more elaborate nervous system. To test if we can detect BMP signaling in the medusozoan nervous system, we performed pSMAD1/5 stainings on the medusa stage of the cubozoan jellyfish *Tripedalia cystophora* (**Fig 7**) and the early juvenile medusa – the so-called ephyra – of the scyphozoan jellyfish *Aurelia coerulea* (**Fig 8**). In *Tripedalia*, we performed co-stainings against pSMAD1/5 and α-Tubulin (αTub), the latter being known to label a large portion of the *Tripedalia* nerve net (Garm et al., 2007). We found that α-Tubulin indeed marked large parts of the *Tripedalia* nerve net, including neurons in the umbrella (**Fig 7B-D’**) and the concentrated ganglia of the nerve ring running along the umbrella rim (**Fig 7E-F)**, but no αTub labeling was observed in the neurons located in the rhopalia, which are the Scyphozoa- and Cubozoa-specific sensory organs containing gravity-sensing statocysts and eyes (**Fig 7I-L)**. In the umbrella, BMP signaling was detectable in αTub-positive ganglionic neurons (**Fig 7C**) and cnidocytes (**Fig 7D**). Cnidocytes are a Cnidaria-specific, highly modified neuronal cell lineage of chemosensory stinging cells identifiable by their distinct crescent-shaped nuclei and the presence of a cnidocyst – the capsule containing the stinging thread (**Fig 7D-D’**). pSMAD1/5 staining was especially pronounced in the cells of the nerve ring (**Fig 7A, E-F’**). Within the nerve ring, we find cells with pSMAD1/5-positive and pSMAD1/5-negative nuclei (**Fig 7E-F’**), which suggests that, like in *Nematostella*, nuclear pSMAD1/5 marks only a subset of neurons in *Tripedalia*. The *Tripedalia* medusa possesses four rhopalia, each containing one statocyst with a heavy statolith (sta) and six eyes: a pair of pit eyes (pe), a pair of slit eyes (se), as well as a much more complex upper lens eye and a lower lens eye (ule, lle) (**Fig 7G-H’**). Since the statolith is located off-center and the whole rhopalium is suspended on a flexible stalk, each rhopalium maintains its orientation in relation to the vector of gravity. Thus, each rhopalium has a clear top side and a clear bottom side. pSMAD1/5-positive cells were restricted to the top side of the rhopalium, including the rhopalium stalk (st), the cells at the stalk base and towards the upper lens eye (**Fig 7J-L**). In the regions of the upper lens eye and pit eye, pSMAD1/5 activity is limited to cells closest to the stalk region. While neurons of the rhopalia are not marked by αTub, many pSMAD1/5-positive cells in the area between the stalk and the upper lens eye are likely to be neurons based on their location and cell morphology. pSMAD1/5-positive cells in this area exhibit a high cytoplasm to nucleus ratio (**Fig 7J, white arrowheads**), resembling the cell morphology of the giant neurons close to the stalk or the balloon cells close to the upper lens eye (Nielsen et al., 2021; Skogh et al., 2006).

**Fig 7.**
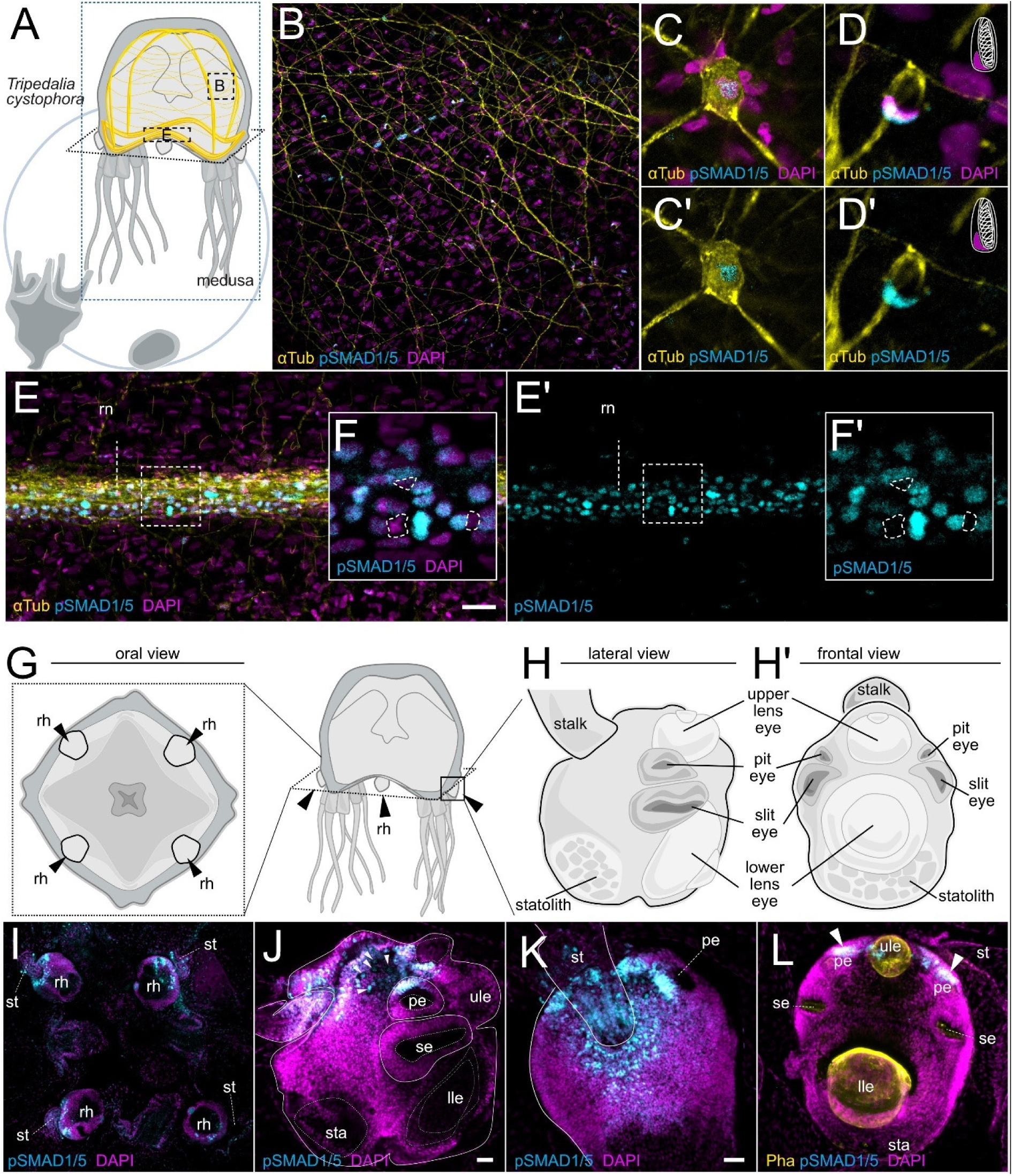
In the box jellyfish *Tripedalia*, BMP signaling is active in ganglionic neurons of the umbrella, the ring nerve and in the upper-most regions of the rhopalium. (A) Life cyle schematic of *Tripedalia cystophora*. (B) α-tubulin antibody staining highlighting the nerve net in the umbrella. (C-C’) detail of alpha-tubulin/pSMAD1/5 double-positive neuron in the umbrella. (D-D’) detail of α-tubulin/pSMAD1/5 double-positive cnidocyte the umbrella. Note the crescent-shaped nucleus at the periphery of the unstained cnidocyst. (E) pronounced α-Tubulin and pSMAD1/5 staining of the ring nerve (F-F’) detail of pSMAD1/5-positive and -negative cells in the ring nerve. (G) Oral view of the umbrella, black arrowheads highlight the location of the rhopalia (rh). (H) lateral and (H’) frontal detail of the rhopalium anatomy. (I) pSMAD1/5 activity in the rhopalia. (J) lateral view of the rhopalium, pSMAD1/5 activity is restricted to the area between the stalk and the pit eye. (K) View from the top highlights pSMAD1/5 activity around the stalk. (L) Frontal view, arrowheads highlight the pSMAD1/5 activity at the two pit eyes. rn - ring nerve, rh - rhopalium, st - stalk, pe - pit eye, se - slit eye, ule - upper lens eye, lle - lower lens eye, sta - statolith; Scale bar 25 µm.

**Fig 8.**
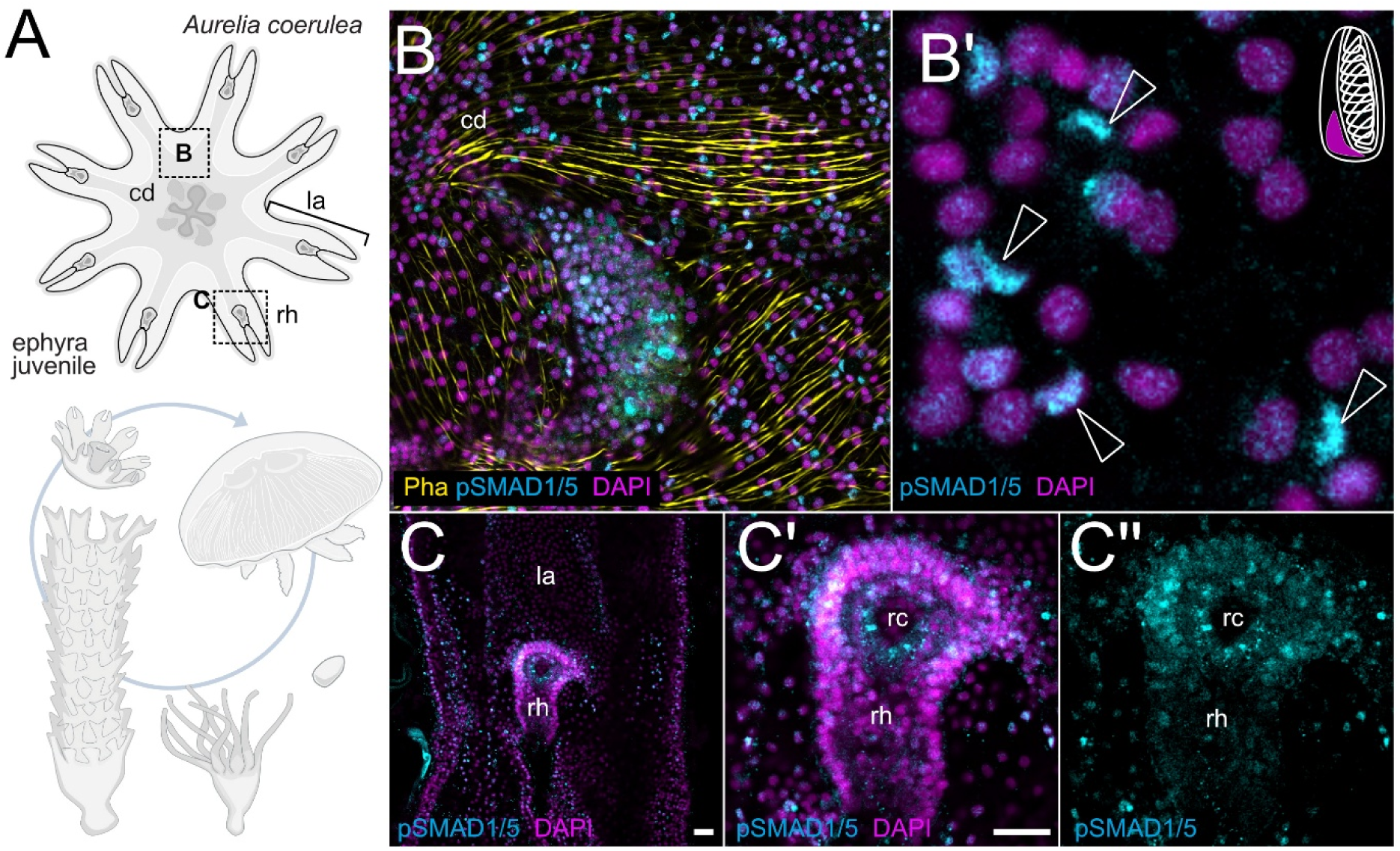
BMP signaling activity in the ephyra of the moon jellyfish *Aurelia* is present in the cnidocytes and parts of the rhopalia. (A) Life cyle schematic of *Aurelia coerulea*, details of the ephyra juvenile stage. (B) pSMAD1/5 staining in the central disc of the ephyra. (B’) Detail showing pSMAD1/5-positive crescent-shaped nuclei typical for cnidocytes. (C) Rhopalia in the lappets exhibit pSMAD1/5 staining at the rhopalic canal (rc). cd - central disc, rh - rhopalium, rc - rhopalic canal; Scale bar 25 µm.

In the *Aurelia* ephyra, we were unable to combine the staining of neuronal markers with pSMAD1/5 staining, which prevented us from detecting the neuronal cells. However, we again observed pSMAD1/5-positive cnidocytes in different parts of the central disc (**Fig 8**). BMP signaling was also detectable in the rhopalia of the ephyra, which are located at the distal end of each of the eight lappets. In the rhopalia, pSMAD1/5-positive cells were mostly located around the rhopalic canal (rc) (**Fig 8**).

### BMP signaling is active in the neurogenic domain of a spiralian

Arrow worms (Chaetognatha) are planktonic predators named after their dart-shaped body and impressive jaw apparatus. Phylogenomic analyses place them within or as a sister group to Gnathifera, within the superphylum Spiralia (Laumer et al., 2019; Marletaz et al., 2019). Chaetognaths possess a brain composed of several ganglia and a centralized ventral nervous system (ventral nerve center), both of which develop paired ventrolateral neuroectodermal cell clusters during embryogenesis (Ordoñez & Wollesen, 2024). In line with observations from other protostomes, we find dorsal expression of *bmp2/4* and ventral expression of *chordin* in the mid gastrula of *Spadella cephaloptera* **(S1 File - Fig. 2)**. Antibody staining shows a dorsal-to-ventral gradient of nuclear pSMAD1/5 in gastrula and early elongation stages (**Fig 9A-C’**). Importantly, pSMAD1/5 signal is observed in the neuroectodermal regions that later contribute to lateral somata clusters of the ventral nerve center (**Fig 9B-C’**). In the hatchling and juvenile, the pSMAD1/5 staining shows broader distribution, including in mesodermal and epidermal derivatives. Nonetheless, BMP signaling remains prominent in the lateral somata, as well as in a number of dorsal sensory organs such as the *pax6*-expressing corona ciliata in the head and ciliary sensory organs of the trunk (**Fig 9D-E’,** (Ordoñez & Wollesen, 2024)). These results show that parts of the central nervous system of *Spadella*, particularly the dorsal regions of the neuroectoderm, are located within the pSMAD1/5-positive domain, and that BMP signaling remains active in the ventral nerve center and in the forming sensory organs later in development.

**Fig 9.**
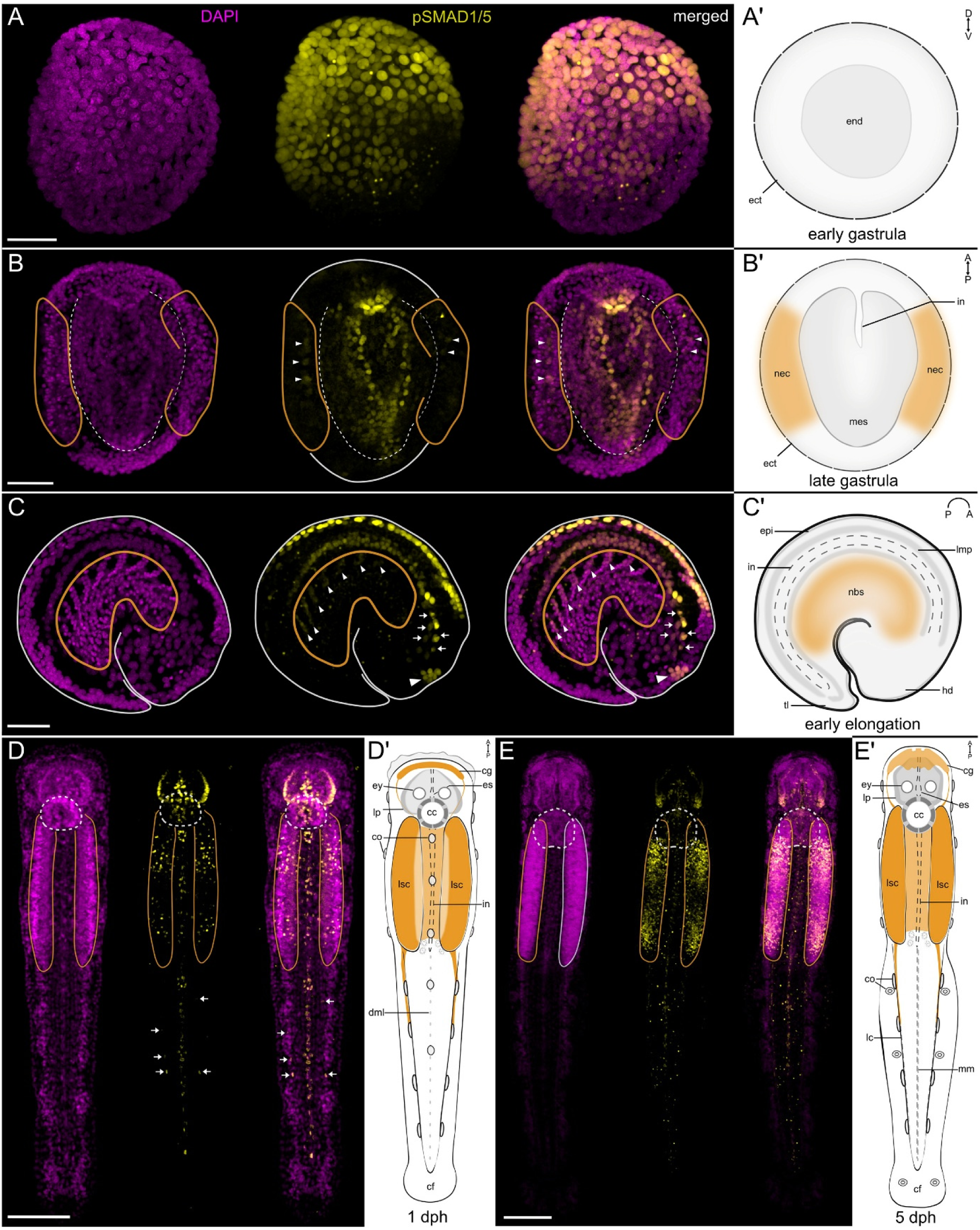
BMP signaling is active in the developing nervous system of the chaetognathe *Spadella*. (A-A’) Transverse view of an early gastrula (A), and schematic representation of its anatomy (A’). Nuclear pSMAD1/5 forms a dorsal-to-ventral gradient. pSMAD1/5-positive nuclei are observed in the dorsal ectoderm and endoderm (ect, end on A’). (B-B’) Dorsal view of a late gastrula (B), and schematic representation of its anatomy (B’). Nuclear pSMAD1/5 is detected in the dorsal mesoderm (mes on B’) and in some neuroectodermal nuclei (arrowheads; nec on B’). (C-C’) Lateral view of the early elongation stage embryo (C) and schematic representation of its anatomy (C’). pSMAD1/5-positive nuclei are present in the dorsal epidermis (epi on C’), the developing digestive tract (in on C’), including the presumptive foregut (arrows), dorsal neuroblasts (white arrowheads) of the developing ventral nerve center (nbs on C’), and presumptive progenitor cells of the inner corona ciliata (red arrowheads). (D, D’) Dorsal view of the *Spadella* hatchling (D) and schematic representation of its anatomy (D’). (D) In the head of the hatchling, pSMAD1/5-positive nuclei are detected in the esophagus (es on D’), lateral plate (lp on D’). Additional signal is observed in sensory structures: corona ciliata (inner cells; cc on D’) and dorsal ciliary tuft/fence organs (co on D’). pSMAD1/5-positive nuclei are also present in the dorsal epidermal midline extending along the trunk and tail (dml on D’), the intestine (in on D’), and lateral somata clusters of the ventral nerve center (lsc on D’). Some pSMAD1/5-positive nuclei of unknown identity are also located lateral to the tail midline (arrows). (E, E’) Dorsal view of the juvenile (E) and schematic representation of its anatomy (E’). pSMAD1/5-positive nuclei remain present in the esophagus, lateral plate, and intestine (es, lp, in on E’), and are particularly prominent in the lateral somata clusters of the ventral nerve center (lsc on E’). Expression is also detected in mesoderm-derived lateral cells (lc on E’) and mesenterial cells (mm on E’). pSMAD1/5 signal is shown in yellow, nuclei are stained with DAPI (magenta). Orange outlines demarcate the neuroectoderm (B) and the developing ventral nerve center(C–E). The dashed outline in (B) marks the developing mesoderm and intestine. Dashed circles in (D) and (E) indicate the corona ciliata. Scale bars, 50 µm (A - C) and 100 µm (D, E). Abbreviations: cc, corona ciliata; cf, caudal fin; cg, cerebral ganglion; co, ciliary tuft/fence organs; dml, dorsal epidermal midline cells; ect, ectoderm; end, endoderm; epi, epidermis; es, esophagus; ey, eye; hd, head; in, intestine; lc, lateral cells; lmp, longitudinal muscle cell precursors; lp, lateral plate cells; lsc, lateral somata cluster; mes, mesoderm; mm, mesenterial cells; nbs, neuroblast of the developing VNC; nec, neuroectoderm; tl, tail bud.

## Discussion

### BMP signaling promotes neural gene expression in *Nematostella*

Regulatory functions of BMP signaling are manifold in Bilateria, and this multifunctionality also extends to their closest relatives, the cnidarians (Knabl et al., 2025). In this study, we analyzed the activity of BMP signaling in a part of the gastrodermal nervous system of the sea anemone *Nematostella.* The overall neural architecture in the polyp has been described before, presenting multiple types of sensory and ganglion neurons with stereotypical placement in the body column (Havrilak et al., 2017; Marlow et al., 2009; Nakanishi et al., 2012; Rentzsch et al., 2017). Different neuron types are present in the two body layers, the epidermis and the gastrodermis, and can be generated by the respective tissue layer (Nakanishi et al., 2012). In the gastrodermis of the polyp, subsets of neurons are regionalized and accumulate in proximity to the longitudinal musculature in the mesentery, which is especially pronounced in the longitudinal neurite tracts along the parietal muscle (Havrilak et al., 2017; Nakanishi et al., 2012). In this region of neural condensations, we observed pronounced BMP signaling activity, partially overlapping with specific subsets of neurons and neuronal progenitors (**Fig 10A**). We found elevated activity of BMP signaling in neurons of the gastrodermis, specifically within the retractor and the parietal muscle regions. BMP signaling was overlapping with neurons expressing *elav1* and *nanos1*, but it was absent from other neurons located in the same regions, *e.g.* expressing *prdm14d*. BMP signaling was also present in two types of *soxb(2)*-positive neurons, firstly in neurons with differentiated morphology in the epithelium and secondly, in basiepithelial cells, likely representing neuronal progenitor cells, based on their localization in the mesoglea. In line with that, analysis of the single-cell RNA-Seq dataset shows the expression of BMP signaling components in the pSC.NPC.1 subset of putative stem cells/neuronal progenitor cells (**S1 File – Fig. 3**). Notably, the occurrence of pSMAD1/5-positive basiepithelial cells, which were also *soxb(2)*-positive in most cases and rarely *soxb(2)*-negative, seemed to be restricted to the neuromuscular mesentery region, while basiepithelial cells in other parts of the body were always pSMAD1/5-negative. These findings are especially interesting in light of the recent findings showing that basiepithelial vasa2/piwi1 double-positive cells represent a population of putative multipotent stem cells of the mesentery, which contributes to at least a portion of the neuroglandular/-secretory repertoire of the gastrodermis in the juvenile and adult polyp (Miramon-Puertolas et al., 2024). The progeny of these vasa2/piwi1-positive cells include *soxB(2)*-positive and *prdm14d*-positive neuronal progenitors, as well as *elav1-*positive neural and *insm1*-positive neuroglandular cells (Miramon-Puertolas et al., 2024). Previously, we could determine that basiepithelial vasa2-positive cells located in the medial mesentery and septal filament have no detectable BMP signaling activity, indicating BMP signaling is absent from putative multipotent stem cells (Knabl et al., 2025). In the context of the proposed model for cell lineage relationships in the gastrodermis (Miramon-Puertolas et al., 2024), we can speculate that BMP signaling activity might correspond to specific branches of the neuroglandular lineage (**Fig 10B**). In this context, BMP signaling might be involved in the decision making of neuronal progenitors and their progeny, which is supported by the transcriptional response upon BMP signaling inhibition.

**Fig 10.**
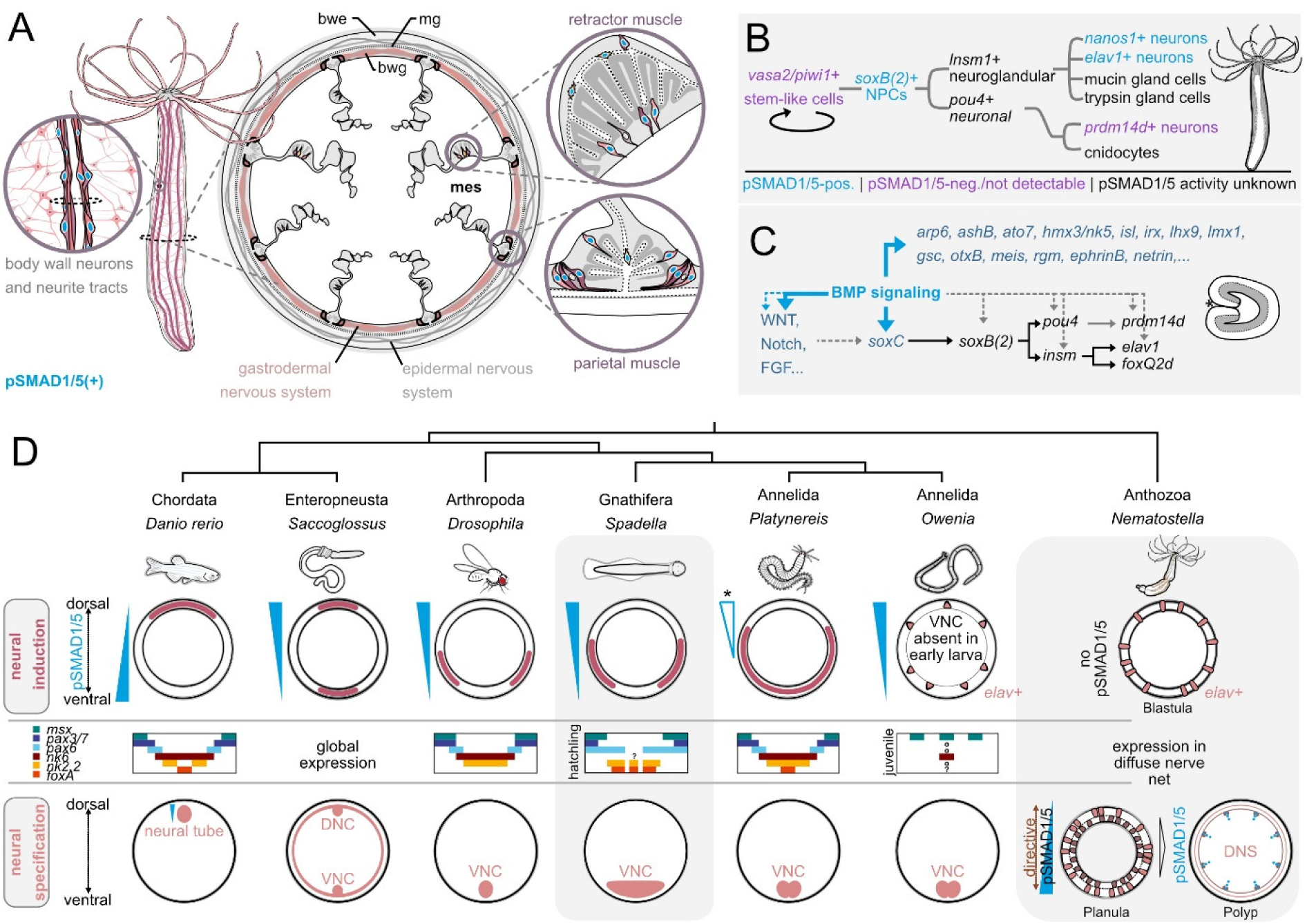
Summary of BMP signaling in the neuronal lineage of *Nematostella* and comparison of neuronal development and graded BMP signaling activity in different Bilateria and *Nematostella*. (A) Localization of pSMAD1/5-positive and -negative neurons in the gastrodermal nervous system. (B) A model for BMP activity during neuronal differentiation in adult polyps. (C) BMP signaling-mediated regulation of neuronal genes in the planula larvae. (D) Comparative overview of phylogenetic relationships, regions of neural induction, expression profiles of mediolateral patterning genes, regions of neural specification and nervous system anatomy in Bilateria and *Nematostella*. bwe-body wall epidermis, mg-mesoglea, bwg-body wall gastrodermis, mes-mesentery, DNC-dorsal nerve cord, VNC-ventral nerve cord, DNS-diffuse nervous system.

Our bulk RNA-seq analysis of K02288 treated adult tissue revealed differential expression of multiple neuronal transcription factors including *soxB(2)*, *irx*, *ap2, ashB* and *paxB* (Layden et al., 2012; Matus et al., 2007; Richards & Rentzsch, 2015; Sinigaglia et al., 2013). Several gene targets in the transcriptomic data from the adult also overlapped with target genes of our previously published RNA-seq data and anti-pSMAD1/5 ChIP data from early stages of development (Knabl et al., 2024). In the datasets from the early developmental stages, we found that many neuronal genes were differentially regulated in the 2d planula upon BMP signaling inhibition by BMP2/4 morpholino-mediated knockdown (bmp2/4MO) and/or direct gene targets of BMP signaling at Late gastrula and 4d planula stage. Based on the overlap of these datasets, BMP signaling appears to regulate neuronal genes both directly and indirectly across different stages of *Nematostella* development (**Fig 10C**). These findings are in agreement with previous analyses of the embryo and the planula larvae suggesting that BMP signaling is involved in patterning and maintenance of the planula nervous system (Watanabe et al., 2014). Using BMP2/4 morpholino knockdown and reverse transcription (RT)-PCR, they described the downregulation of genes encoding for a group of achaete-scute and atonal-related basic helix-loop-helix (bHLH) proteins and of the neuropeptides *rf-amide* and *glw-amide* (Watanabe et al., 2014). Our RNA-seq analysis of bmp2/4MO planulae yielded similar results, expanding our list of neurogenic factors, the majority of which proved to be downregulated in the absence of BMP signaling. Downregulated genes include *soxb(2)*, marking neuronal progenitors that give rise to nematocytes, sensory and ganglion neurons (Richards & Rentzsch, 2014, 2015), as well as *soxC*, an upstream regulator of *soxB(2)* (Steger et al., 2022), which we have previously identified as a direct gene target of pSMAD1/5 (Knabl et al., 2024). Moreover, among the downregulated genes we find markers of neural progenitors and neural differentiation (*pou4, insm, elav1, prdm14d, nanos1, otp;* (Lemaitre et al., 2023; Nakamura et al., 2023; Nakanishi et al., 2012; Tourniere et al., 2020; Tourniere et al., 2022)), bHLH family members *(ashA, ashB, ashC, ashD, arp6, ato7;* (Layden et al., 2012; Richards & Rentzsch, 2015; Watanabe et al., 2014)) and transcription factors associated with neurogenesis in bilaterians (*gata, soxE1, ap2, irx, isl1, pou3, runx;* (Galliot et al., 2009; Rentzsch et al., 2017)). In accordance, RNA detection by in situ hybridization in 2d planula upon BMP2/4 morpholino knockdown or K02288 BMP receptor inhibition showed significant downregulation of neurogenic factors. Notably, low levels of marker gene expression were still detectable, indicating that neurogenesis was not completely abolished, supporting the idea that BMP signaling is not required for the initiation of neurogenesis in the early embryo. Taken together, our data suggest a predominantly pro-neural role of BMP signaling in *Nematostella* larva and adult.

### BMP signaling activity overlaps with neuronal populations in Medusozoa

To determine if BMP signaling also plays a role in the diffuse nervous system of medusozoans, we analyzed the activity of BMP in the box jellyfish *Tripedalia* and the ephyra stage of the moon jellyfish *Aurelia*. Co-stainings of pSMAD1/5 and α-tubulin in *Tripedalia* revealed BMP signaling activity in different parts of the diffuse nerve net of the medusa, highlighting ganglion neurons in the umbrella and the ring nerve. Similar to the situation in *Nematostella*, where we observed that pSMAD1/5 activity differs between *elav*1-positive and *prdm14d*-positive neurons, we found both pSMAD1/5-positive and -negative neurons in the ring nerve of *Tripedalia,* indicating the presence of distinct neuronal subsets. In the ring nerve, the presence of specific neuronal populations labeled by different antibody combinations was demonstrated in the hydromedusa *Cladonema* (Koizumi et al., 2015) and in *Tripedalia* (Garm et al., 2007). In *Tripedalia* and *Aurelia* medusae/ephyrae, we detected BMP signaling activity in distinct parts of the light and gravity sensing rhopalia. The rhopalic nerve net features distinct subsets of regionalized sensory and ganglion cells (Nakanishi et al., 2009; Nielsen et al., 2021; Parkefelt & Ekstrom, 2009; Skogh et al., 2006). In *Aurelia*, BMP signaling activity was more pronounced around the rhopalic canal compared to the more distal intermediate segment. The regionalization of BMP signaling was even more pronounced in the rhopalia of *Tripedalia*, where areas of pSMAD1/5 activity were restricted to the region adjacent to the rhopalium stalk. pSMAD1/5 activity is found in the areas known to be populated by specific neurons, including PCNA-positive neurons innervating the pit eye and the upper lens eye, the giant neurons or ballon cells and FMRF-positive neurons around the stalk, but not flank neurons or retina-associated neurons (**Suppl. Fig**, (Skogh et al., 2006)). In both *Tripedlia* and *Aurelia*, we also identified pSMAD1/5 activity in the stinging cells (cnidocytes), a highly specialized cell type of the neurosecretory lineage across Cnidaria (Galliot et al., 2009; Steger et al., 2022). In contrast, in anthozoan *Nematostella*, the expression of BMP pathway genes is upregulated in mature cnidocytes according to single-cell data analysis, but we never detected nuclear pSMAD1/5 staining in any type of stinging cell (Knabl et al., 2025). In summary, both in medusozoan and anthozoan cnidarians, we detect neurons receiving BMP signals, which suggests that pro-neural role of BMP signaling is an ancestral cnidarian trait predating the anthozoan-medusozoan divergence.

### BMP signaling promotes neurogenesis in parts of bilaterian nervous systems

BMP signaling is associated with various stages of neurogenesis in Bilateria, displaying negative or positive regulatory effects depending on the specific location and timing within the developmental context. In vertebrates and arthropods, the specification of the centralized nervous system requires inhibition of BMP signaling, while differing levels of BMP activity regulate the mediolateral patterning of neurectoderm by homeobox factors *nk2.1, nk2.2, nk6, pax6, pax3/7* and *msx* (Holland et al., 2013) (**Fig 10D**). The question about neurogenic functions of BMP signaling becomes more complicated when we consider animal groups outside of vertebrates and arthropods. Spiralians, ambulacrarians and xenacoelomorphs display an assortment of centralized and decentralized neural architectures that, in many cases, differ in their developmental regulation from what is known from groups with centralized nervous systems.

Little is known about neuronal roles of BMP signaling in xenacoelomorphs, whose phylogenetic position, either at the base of Bilateria or within the deuterostomes, is still unresolved (Alvarez-Presas et al., 2024; Cannon et al., 2016; Kapli et al., 2021; Martinez et al., 2024; Philippe et al., 2011; Schiffer et al., 2024). The group displays divergent neural architectures, ranging from diffuse nets with simple nerve plexuses to organized nervous systems with multiple nerve cords (Alvarez-Presas et al., 2024). BMP signaling inhibition in a xenacoelomorph nemertodermatid did not affect the establishment of the nerve cords but increased the number of serotonin/(5-HT)-positive commissures (Martin-Duran et al., 2018).

In non-chordate deuterostomes, different forms of decentralized nervous systems exist, including the pentaradial nerve cords and nerve ring in echinoderms and the epidermal nerve net in hemichordates, also featuring a ventral and the dorsal nerve cord (Adameyko, 2023). Like in chordates, BMP signaling appears to initially repress neurogenesis in ambularcrarians (Su et al., 2019), and later, may be required for local neural specification (Hinman & Burke, 2018). In the sea urchin, BMP signaling locally promotes the proneural markers *soxC*, *soxB1* and *sip1,* as well as serotonergic neurogenesis only in the anterior neural ectoderm, but not in the ciliary band or in the region of the endomesoderm that later will give rise to the gut (McClay et al., 2018). In contrast, BMP signaling disruption in the sea star had no effect on *elav* expression but affected the localization of *elav*-positive cells (Yankura et al., 2013). In the hemichordate *Saccoglossus*, exogenous BMP had no anti-neurogenic effects, based on the persisting expression of pan-neural marker *elav* (Lowe et al., 2006).

Neural anatomies in spiralians are diverse, often displaying different numbers of nerve chords. In the context of neurogenesis, BMP signaling was found to have positive or negative effects on the formation or organization of nerve chords, the brain or eyes (Lambert et al., 2016; Lanza & Seaver, 2020a, 2020b; Tan et al., 2022; Webster et al., 2021; Webster & Meyer, 2024). In the snail *Ilyanassa*, BMP signaling was shown to promote the neural ectoderm and the formation of eyes and the shell (Lambert et al., 2016), while another gastropod, *Crepidula*, exhibits enlarged cerebral ganglia and ectopic ocelli when BMP signaling is blocked by the selective Activin receptor-like kinase-2 inhibitor Dorsomorphin Homolog 1 (DMH1) (Lyons et al., 2020). In the annelid worm *Chaetopterus*, BMP signaling inhibition had no effect on neural structures (Lanza & Seaver, 2020a). Similarly, a recent analysis in the early branching annelid *Owenia*, where a BMP signaling gradient is responsible for the DV patterning, showed that modulating BMP signaling also did not prevent the development of neural structures or differentiation of specific neuron types (Carrillo-Baltodano et al., 2025). Until now, no information on the areas of BMP signaling activity was available for the non-lophotrochozoan spiralians. We analyzed BMP signaling activity in the embryos, hatchlings and juveniles of the chaetognath *Spadella* and showed that a dorsal-to-ventral BMP signaling gradient forms during its early development. *Spadella* has a highly centralized ventral nervous system, the ventral nerve center, which appears to be patterned mediolaterally by the same genes as the central nervous systems of vertebrates, insects and polychaetes (Ordoñez & Wollesen, 2024). Importantly, the chaetognath ventral nerve center forms from two bilateral patches of neurectoderm located in the ventrolateral sides of the ectoderm of the embryo. These regions give rise to multiple neural components, including the lateral somata clusters, which contribute to the developing nervous system. Here we show that during *Spadella* development, BMP signaling is active in the dorsal parts of neurectoderm of late gastrula and subsequently in the dorsal parts of the lateral somata clusters of early elongation stage embryos. Later, in the hatchling and in the juvenile, BMP signaling is strong in parts of the ventral nerve center and transiently also in some of the sensory organs such as the corona ciliata.

Chaetognaths are not the only animals with highly centralized nervous systems, where BMP signaling is active in the neural structures – especially in the peripheral and enteric neurons. In the annelid worm *Platynereis* and the fly *Drosophila*, which both have a ventral nerve cord, the expression of pro-neural marker *atonal,* as well as the formation of peripheral sensory neurons were promoted by BMP signaling (Denes et al., 2007; Rusten et al., 2002). In the cephalochordate *Amphioxus*, BMP signaling inhibition is required for CNS formation (Onai et al., 2010), whereas high BMP signaling promotes the formation of *elav*-positive peripheral sensory neurons (Lu et al., 2012). Similarly, BMP signaling in the tunicate *Ciona* is involved in the induction and differentiation of the peripheral nervous system (Pasini et al., 2006; Waki et al., 2015). In several vertebrate models (fish, chick, mouse and human), BMP signaling is associated with promoting the enteric nervous system (Chalazonitis et al., 2008; Moore et al., 2024). In the zebrafish, pSMAD1/5 activity overlaps with *phox2b*-positive enteric progenitor cells and *elav*-positive differentiated enteric neurons, regulating either the generation of progenitor cells or their differentiation at different time points of embryonic and larval development (Moore et al., 2024). Taken together, rather than being an unusual feature of cnidarians with their simple, diffuse nervous systems, pro-neural BMP signaling activity is also observed across Bilateria where it promotes neurogenesis of peripheral, sensory and enteric neurons.

Our analyses suggest that positive neuronal regulation by BMP signaling may represent an ancestral feature, predating the cnidarian-bilaterian divergence. Therefore, we propose that the “anti-neural” role of BMP signaling documented in vertebrates and arthropods (**Fig 10D**) is a result of the co-option of the global program of the BMP-dependent DV patterning of the ectoderm in the animals where neurectoderm is represented by a contiguous domain at the “low BMP signaling” side of the DV axis. In the future, it will be important to analyze the expression and the functions of the transcription factors responsible for the mediolateral patterning of the central nervous system in bilaterians with multiple nerve cords as well as in the ones with diffuse nervous systems. If we find evidence of involvement of the mediolateral patterning transcription factors in the global patterning of the ectoderm rather than in the neural patterning, this will be a strong argument in favor of the “simple urbilaterian” hypothesis (Hejnol & Martindale, 2008) and multiple independent centralizations of the nervous system across the bilaterian tree.

## Materials and methods

### Animal culture

Adult polyps of *Nematostella vectensis* adult polyps were maintained in the dark at 18°C in 16 ppm artificial seawater (*Nematostella* medium, NM) and spawned as described before (Fritzenwanker & Technau, 2002; Genikhovich & Technau, 2009). *Tripedalia* medusae were collected at the aquarium of the Tiergarten Schönbrunn in Vienna and fixed and processed for antibody staining on the same day. *Aurelia* polyps were kept in petri dishes in artificial sea water (ASW, 35‰) at 20°C and fed with *Artemia* once a week. Strobilation was induced by keeping the polyps at 15°C over night. Ephyra were transferred to a small beaker with ASW and fed with *Artemia* once a week. Live *Spadella cephaloptera* specimens were collected from the intertidal zone near Roscoff, France, and maintained under laboratory conditions until spawning, following established procedures (Ordoñez & Wollesen, 2024; Wollesen et al., 2023).

### Single-cell RNA-seq analysis

Analysis of single-cell transcriptomic data was performed using previously published data (Cole et al., 2024) (https://github.com/technau). Gene expression of genes of interest in the developmental dataset and the neuroglandular subset was examined using the Seurat::DotPlot function with maximum dot size fixed at 100%. All R scripts used for the processing and analysis of the *Nematostella* single-cell data are available on GitHub (https://github.com/technau/Nv2_Atlas).

### Immunostaining and vibratome sectioning

Antibody staining in *Nematostella* polyps, *Aurelia* ephyra and *Tripedalia* medusa was performed as described previously (Knabl et al., 2025). Samples were incubated in Blocking solution (1% BSA, 5% sheep serum, 1 × PBS, 0.2% Triton-X100, 20% DMSO) at room temperature for at least 2 h. The primary antibodies (1:200 rabbit anti-pSMAD1/5/9 (mAb #13,820, Cell signaling)) were pre-absorbed in Blocking solution (1% BSA, 5% sheep serum, 1 × PBS, 0.2% Triton-X100, 0.1% DMSO) and incubated with the sample at 4 °C overnight. Samples were washed ten times in 1 × PBS / 0.2% Triton-X100, incubated in Blocking solution at room temperature for 1 h and stained with the secondary antibodies at room temperature for 2 h or at 4 °C overnight. Secondary antibodies were diluted 1:1000 in Blocking solution (goat α-rabbit IgG-Alexa633 (Invitrogen A21070), goat α-mouse IgG-Alexa488 (Invitrogen A11001), goat α-mouse IgG-Alexa568 (Invitrogen A11004) and DAPI, 1:1000) Samples were washed ten times in 1 × PBS / 0.2% Triton-X100, and either infiltrated with and mounted in VECTASHIELD® Antifade Mounting Medium (H-1000–10, VectorLabs) or processed for vibratome sectioning. Samples were embedded in 10% gelatin/PBS, fixed in 4% Formaldehyde in PBS at 4 °C overnight and sectioned at a Leica VT1200 vibratome as described before. All stainings were performed at least two times independently with three or more animals imaged. Images processing and figure preparation were performed using Fiji (Schindelin et al., 2012) and Affinity Designer (https://affinity.serif.com/en-us/designer/).

*Spadella* embryos and post-hatching stages were fixed in 4% paraformaldehyde in MOPS buffer (0.1 M MOPS pH 7.4, 2 mM EGTA, 1 mM MgSO_4_, 2.5M NaCl). For pSMAD1/5 immunostaining, samples were stored in 70% ethanol at −20°C. pSMAD1/5 immunostaining was performed on early and late gastrulae, early elongation stage individuals, hatchlings, and early juveniles (five days post-hatching). Samples were rehydrated through an ethanol/PBS series (50%, 25%, PBS) and washed three times in PBS-TX (0.3% Triton X-100 in PBS). Blocking was carried out overnight at 4°C in PBS-TX with 3% normal goat serum (NGS), followed by overnight incubation with anti-pSMAD1/5/9 antibody (1:100; Cell Signaling #13820) in PBS-TX + 3% NGS. After a series of washes in PBS-TX for two hours, specimens were incubated overnight at 4°C with goat anti-rabbit Alexa Fluor 488 (1:1000; Invitrogen #A-11094) and DAPI (1 μg/mL) in PBS-TX + 1% NGS. After a final series of washes, samples were mounted in Vectashield® Antifade Mounting Medium (Vector Laboratories).

### Morpholino knockdowns, inhibitor treatments and in situ hybridization

For knockdowns of *bmp2/4* and *gdf5-like* in *Nematostella*, morpholino oligos were injected into fertilized eggs as described before (Genikhovich et al., 2015; Saina et al., 2009). For chemical BMP signaling inhibition, animals were incubated in *Nematostella* Medium (NM) with 6 µM K02288 or, as control treatment, with equal volumes of DMSO. Treatments were carried out starting from blastula stage (18 hours post fertilization, hpf) For the wash-out at 2-day (48 hpf), 4-day (96 hpf) or 5-day (120 hpf) planula stage, animals were washed three times with NM. Morpholino and K02288 treated animals were fixed for In situ hybridization as described before (Knabl et al., 2024; Lebedeva et al., 2021) with minor modifications. *Nematostella* planulae were fixed in 4% PFA in 1× PBS and 0.1% Tween 20 (PTW) for 1 hr at room temperature. For permeabilization, samples were incubated in 10 μg/ml Proteinase K/PTW for 20 min at room temperature. Following the 2× SSC wash, 4d planulae were incubated in 1 unit/μl RNAseT1 in 2× SSC at 37°C for 40 min and then washed with 0.075× SSC to reduce unspecific “collar” staining around the pharynx.

*Spadella* embryos used for HCR-FISH with *bmp2/4* and *chordin* probes were fixed at gastrula stage as described above and stored in 100% methanol at −20°C (rather than in 70% ethanol). HCR probes were designed with the *insitu_probe_generator.py* script (https://github.com/rwnull/insitu_probe_generator; (Kuehn et al., 2022) and synthesized by Integrated DNA Technologies. Hairpins were purchased from Molecular Instruments (USA). HCR-FISH followed the protocol of Choi et al. (Choi et al., 2016), with additional modifications from Bruce et al. (Bruce et al., 2021). Cell nuclei were counterstained with DAPI during the final incubation step. Samples were then incubated in Vectashield® for 20 minutes, cleared in a graded TDE (2,2′-thiodiethanol) series in PBS (30%, 60%, and 80%), and mounted in 80% TDE for imaging. Imaging was performed with a Leica TCS SP5 confocal microscope (Leica Microsystems, Germany). Image processing and figure preparation were carried out using Fiji (Schindelin et al., 2012) and Inkscape (https://inkscape.org).

### Inhibitor treatments and bulk RNA-seq of adult polyps

Adult polyps were incubated in 5mM lidocaine in NM for 5 min or until fully relaxed and transferred to a 6-well plate containing 20µM K02288 in NM or DMSO in NM as control. For a more even exposure, the solution was injected into the body column through the mouth using a 1 ml syringe and a blunt Sterican needle with a bent tip. Animals were treated for 5 hours, relaxed for 10 min using 5mM lidocaine in treatment solution and then transferred to 100% MeOH for dissection. Total RNA extraction was performed using the Zymo Research Quick-RNA Miniprep Kit.

*Nematostella vectensis* genome (Zimmermann et al., 2023), differential expression analysis was performed with DeSeq2 (Love et al., 2014). For the analysis, expression changes with an adjusted p-value of <0.05 were used. Datasets generated in this study were compared with the previously published anti-pSMAD1/5 Chip dataset (late gastrula and 4-day planula stage) and the BMP2/4 morpholino (bmp2/4MO) RNA-seq data (2-day planula) (Knabl et al., 2024). The formerly used NVE gene models were matched to the new NV2 gene annotations. NVE gene models with no clear NV2 ID were not considered for further analysis. NVE-to-NV2 matching reduced the number of gene models used for the comparison, e.g. of 254 NVE models identified as direct targets of BMP signaling by ChIP-Seq (Knabl et al., 2024), 210 had a one-to-one NV2 counterpart.

## Supporting information

Table S1. List of differentially expressed genes upon K02288 treatments of adult polyps

Table S2. Comparison of K02288 RNA-Seq, bmp2/4MO RNA-Seq and LG4dP anti-pSMAD1/5 ChIP-Seq datasets

## Acknowledgments

This research was funded in whole, or in part, by the Austrian Science Fund (FWF) grant (DOI 10.55776/P32705) to G.G. and to T.W. (P34665). P.K. is a recipient of the Dimitrov Fellowship of the Austrian Academy of Sciences (OeAW) and of the Writing-up Fellowship of the Konrad Lorenz Institute for Evolution and Cognition Research (KLI). For the purpose of Open Access, the author has applied a CC BY public copyright license to any Author Accepted Manuscript (AAM) version arising from this submission. We thank Fabian Rentzsch (University of Bergen) for providing *soxb(2)::mOrange* and *prdm14d::gfp* transgenic lines of *Nematostella*, Oliver Link (University of Vienna) for proving *Aurelia* ephyra and Roland Halbauer (Tiergarten Schönbrunn, Vienna) for providing *Tripedalia* jellyfish. For the purpose of Open Access, the author has applied a CC BY public copyright license to any Author Accepted Manuscript (AAM) version arising from this submission. Confocal microscopy was performed at the Core Facility Cell Imaging and Ultrastructure Research, University of Vienna - member of the Vienna Life-Science Instruments (VLSI).

## Data availability

Adult *Nematostella* tissue RNA-Seq data is deposited in NCBI at https://www.ncbi.nlm.nih.gov/bioproject/PRJNA1269470

## Supporting information captions

**S1_File** Supplementary Figures S1-3.

**S1 Table**List of differentially expressed genes upon K02288 treatments of adult polyps

**S2 Table** Comparison of K02288 RNA-Seq, bmp2/4MO RNA-Seq and LG4dP anti-pSMAD1/5 ChIP-Seq datasets

## Supplemantry figures

**S1 File - Fig. 1.**
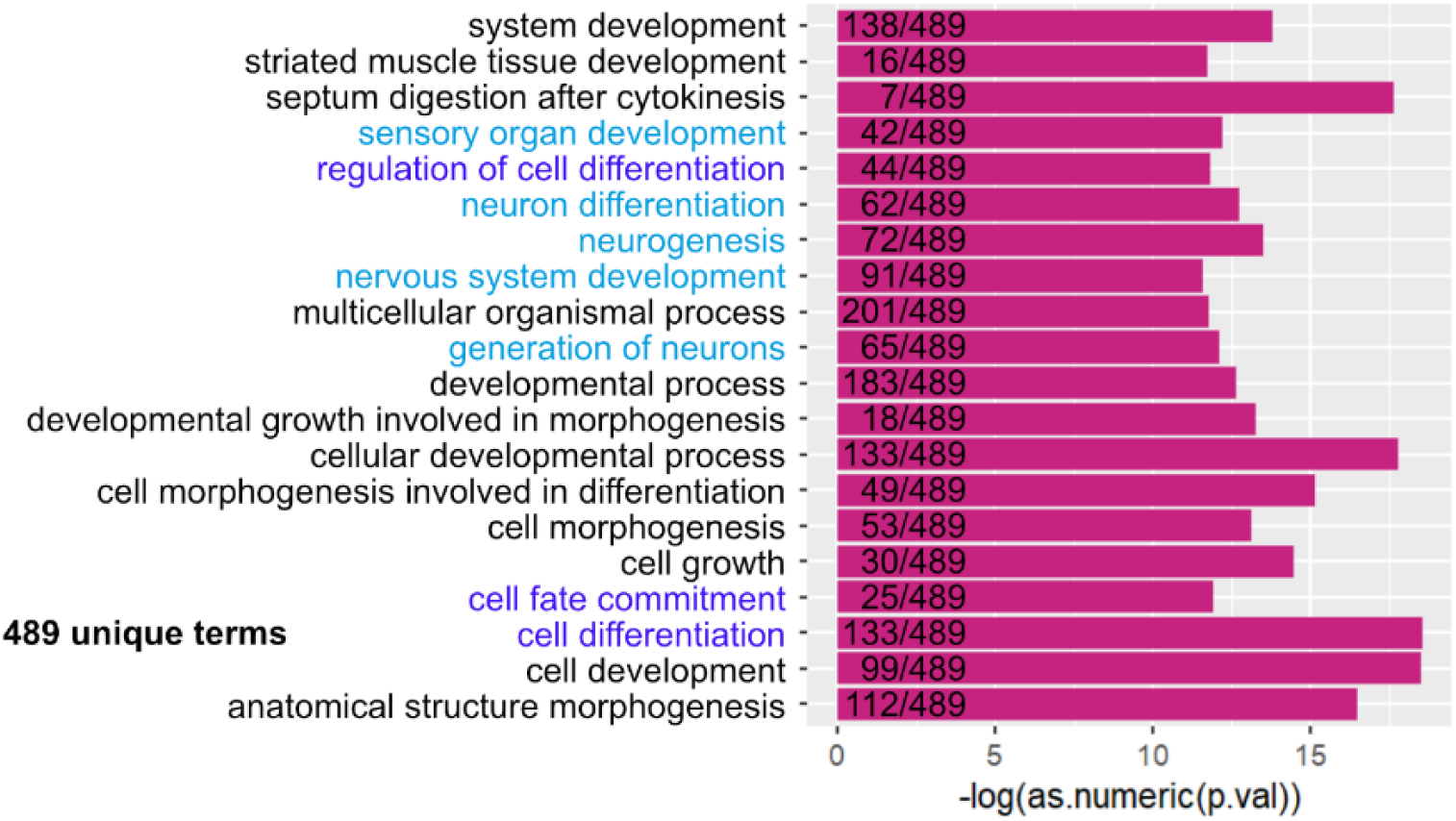
GO-enrichment analysis of the differentially expressed genes upon K02288 treatment of adult polyp tissues.

**S1 File - Fig. 2.**
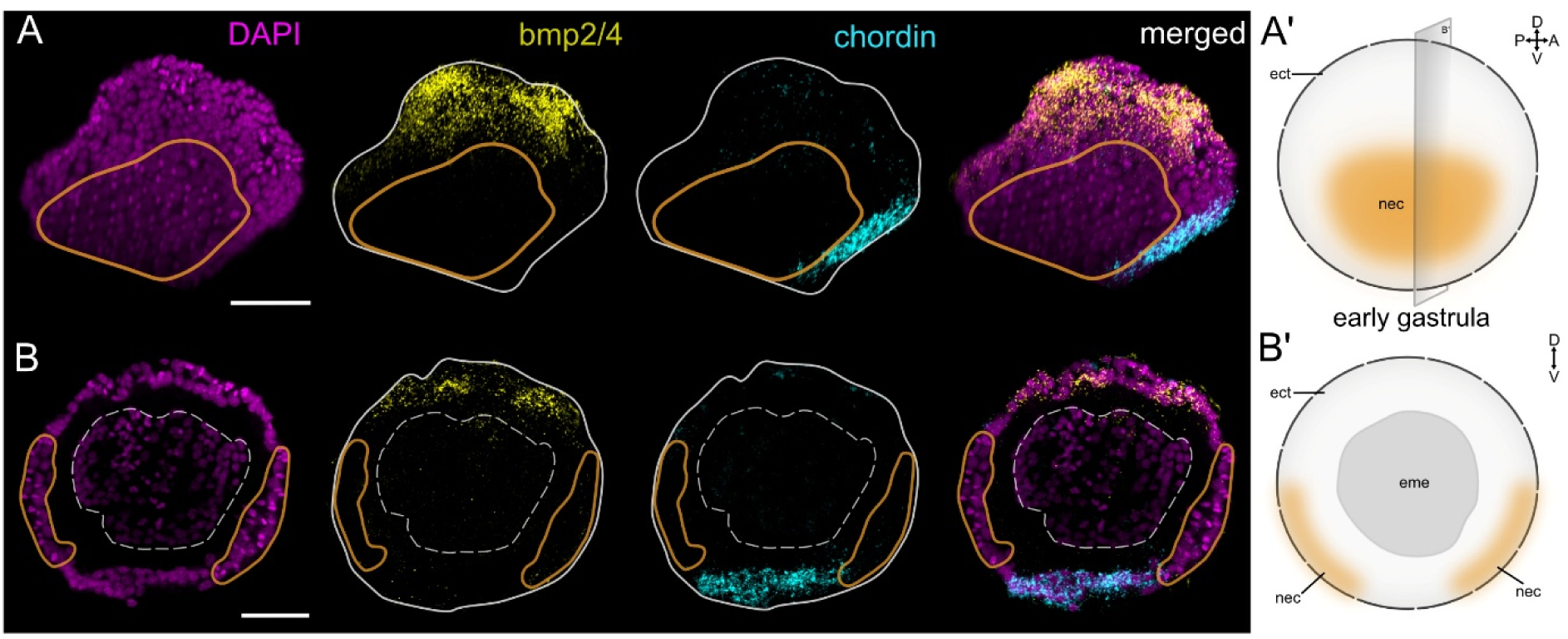
*bmp2/4* is expressed dorsally and *chordin* – ventrally in the mid gastrula of the chaetognath *Spadella*. (A-A’) Lateral view and (B-B’) frontal view and the corresponding sketches of the anatomy of the embryo. Orange outlines demarcate the neuroectoderm (A, B) and dashed outlines demarcate the endoderm (B). Scale bars 50 µm. Abbreviations: ect, ectoderm; eme, endomesoderm; nec, neuroectoderm.

**S1 File - Fig. 3.**
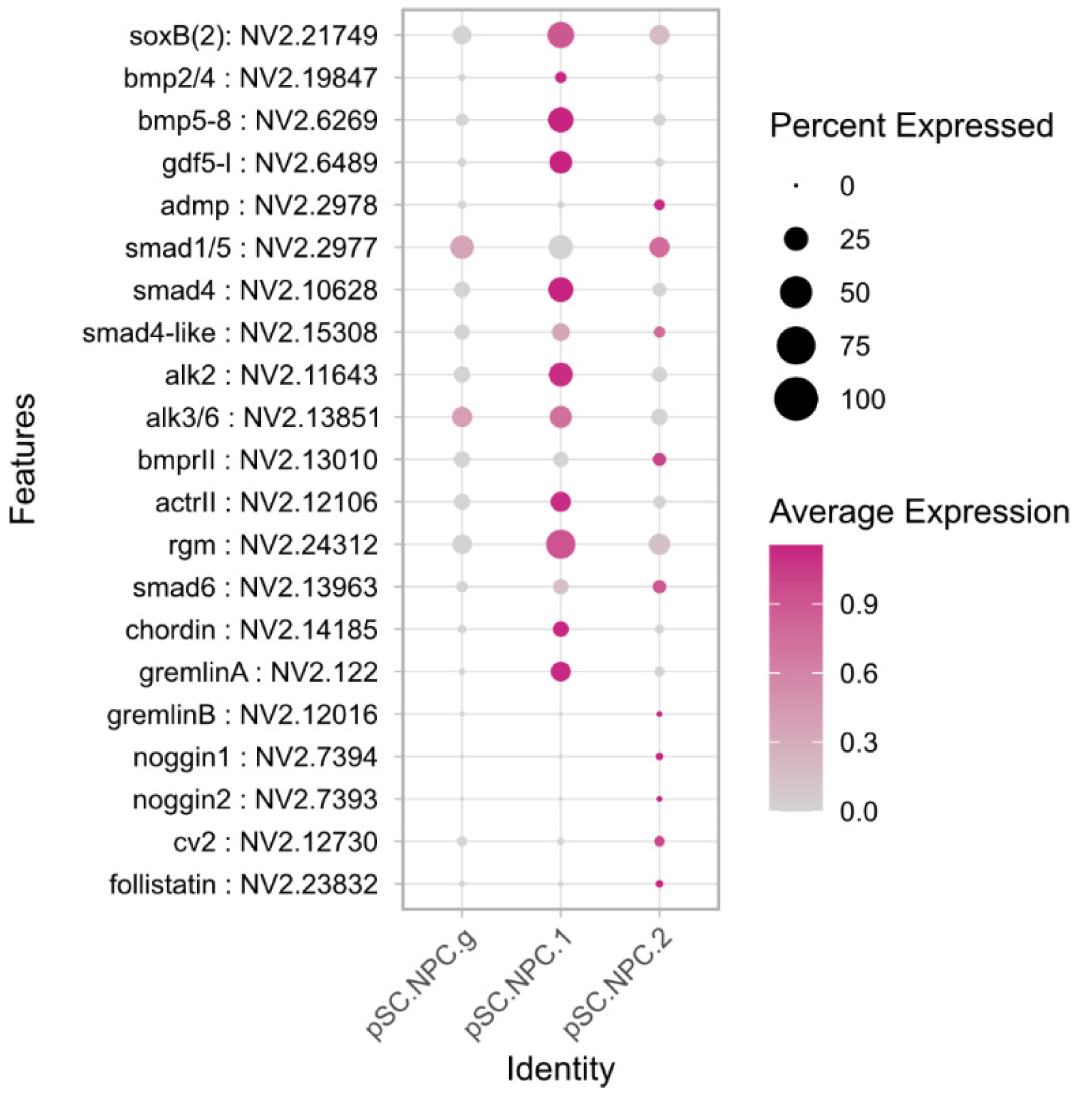
The expression of BMP signaling components in the developmental subset of primordial stem cells and neuronal progenitors is enriched in the *soxB2*-positive subcluster pSC.NPC.1. Average scaled expression of 0 or below is indicated in grey.

